# Individual Differences in the Correspondence Between Psychological and Physiological Stress Indicators

**DOI:** 10.1101/2024.08.23.609328

**Authors:** Kirsten Hilger, Irma Talić, Karl-Heinz Renner

## Abstract

Stress threatens physical and mental health. Reactions to acute stress comprise multiple levels, including negative thoughts, bodily symptoms and behaviors. Individuals differ in their reaction to acute stress, and importantly, also in the extent to which these levels align, with a closer correspondence between psychological and physiological stress indicators being beneficial for mental health and well-being. This preregistered study investigates such individual differences systematically by inducing psychological (social-evaluative) and physiological (cold water) stress with the Maastricht Acute Stress Test (MAST) in 149 healthy adults. Participants indicated their perceived stress and four physiological stress indicators (blood pressure, heart rate, salivary cortisol, alpha-amylase) were obtained. Finally, multiple personality traits were assessed as potential moderators, including the Big Five, trait anxiety, and general cognitive ability. In line with previous research, psychological and physiological stress indicators were only weakly correlated and Bayesian analyses provided evidence favoring the absence of close psychophysiological correspondence. Considering individual differences in personality, especially conscientiousness and openness emerged as potential moderators. We propose individual differences in interoceptive abilities as another critical moderator, which deserves further investigation, and discuss how future research on individual differences in psycho-physiological correspondence can contribute to further our understanding of mental and physical diseases.

## 1. Introduction

Stress has been identified as a major health challenge of the 21^st^ century”[1]. Indeed, numerous studies demonstrate associations between *chronic* stress, i.e., frequent episodes of *acute* stress over a longer period of time without being able to recover, and the onset of physical diseases like hypertension and heart attacks[2,3]. Similarly, for the development of mental disorders like depression, anxiety, and chronic pain, chronic stress was identified as a critical vulnerability factor[4,5]. Large-scale studies further show that chronic stress has substantially increased over the last decades due to working conditions like multitasking, time pressure or the need to adapt to quickly changing technologies[6]. Therefore, it is of pressing importance to understand the mechanisms and peculiarities of acute stress as defining element of chronic stress conditions.

Psychologically, acute stress is characterized by heightened levels of perceived tension and anxiety, influencing cognitive processes such as rumination and intrusive thoughts[7]. Physiological markers of stress encompass alterations in heart rate, blood pressure, skin conductance level and cortisol release[8,9]. Finally, disruptions in sleep patterns, changes in eating habits as well as alcohol and drug consumption exemplify the behavioral manifestations of acute stress[10,11].

From an evolutionary perspective, a correspondence between different response systems of acute stress would be functional in terms of facilitating adequate reactions to external demands[12]. When facing, for instance, a tiger, only a synchronicity between the psychological (quickly perceiving the tiger as a dangerous animal), the physiological (immediate activity of the sympatho-adrenal medullary (SAM) stress axis followed by the hypothalamic-pituitary-adrenal axis (HPA) and the behavioral response system (fight or flight) would ensure survival. Empirical studies, however, only show weak associations between psychological, physiological and behavioral stress indicators with substantial variations across different studies[13,14].

This heterogeneity may be explained by the use of different stress inductions, varying stress measures, and two alternative ways to determine the psycho-physiological correspondence: on a within-subject level, i.e., correspondence of psychological and physiological measures over different trials or across time within individuals[15] versus on a between-subject level, i.e., correspondence of psychological and physiological measures between different persons, so that persons scoring high on psychological measures tend to score also high on physiological measures[16]. With regard to the latter approach, a review of 49 studies revealed that significant correlations between psychological and physiological stress markers were observed in 25% of the studies only. The authors hypothesized that personality traits might be a critical factor moderating the psycho-physiological correspondence but did not test this hypothesis further[13]. Low between-subject psycho-physiological correspondence was also observed in a recent study that used experience sampling and mobile smartwatch sensing to assess psychological and physiological indicators of stress in everyday life[17]. However, these every-day life stress situations were mainly of psychological nature as well. Whether similarly low correspondence between psychological and physiological stress responses exists after facing situations combining psychological and physiological stressors has not yet been investigated.

Personality traits, specifically the Big Five (extraversion, neuroticism, agreeableness, conscientiousness, openness), have been established as both protective and vulnerability factors with regard to individuals’ perception of acute stress. In a recent meta-analysis, neuroticism turned out to be strongly positively, whereas extraversion, agreeableness and conscientiousness were negatively associated with measures of acute psychological stress perception[18]. By contrast, no significant associations were observed between the Big Five and physiological measures of acute stress such as heart rate variability or skin conductance. The *moderating* role of personality traits has been addressed in the well-established Exposure-Reactivity model of stress and coping[19,20], such that e.g., persons high in neuroticism tend to experience stressors more severely than persons low in neuroticism.

Only few studies, however, considered the possible moderating effects of different personality traits or personality-related variables on the psycho-physiological correspondence: A pioneer study examined potential links between psycho-physiological correspondence and attachment style[21], while higher body awareness has been shown to be positively related with psycho-physiological correspondence in another study[16]. Addressing clinical relevance, it has been observed that persons with higher correspondence between self-reported stress and heart rate reported higher psychological well-being, fewer depressive symptoms, and lower levels of trait anxiety, denial coping and proinflammatory biomarkers, altogether supporting the significance of psycho-physiological correspondence for both physical and mental health[14]. Finally, the most recent investigation found higher conscientiousness and neuroticism to be related to higher psycho-physiological correspondence, while extraversion was associated with lower correspondence[17]. Different associations of personality traits with interoceptive abilities (i.e., abilities of perceiving internal bodily signals) were proposed as potential explanation for these findings. Specifically, interoceptive abilities were assumed to be positively associated with conscientiousness[22] and neuroticism[23], while extraversion typically demonstrates no association with interoception, most likely due to the inclination of extraversion towards the outside world[23]. Notably, both studies linking psycho-physiological correspondence to personality factors considered only one single measure as indicator of the physiological stress response. However, the use of a single measure as indicator of the physiological stress system might be problematic, as not all persons react in the same way and there are varying amounts of non-responders in respect to different measures[24,25]. Based on this observation, the use of multiple different physiological indices within the same sample was proposed as a promising means to investigate the link between psycho-physiological correspondence and personality in more detail[14], but this has – to the best of our knowledge – not yet been realized.

The current study provides a comprehensive investigation of the between-subject correspondence between psychological and physiological stress responses in reaction to acute stress. We address the current research gaps (a) by applying a stress induction procedure that comprises exposure to psychological and physiological stressors and causes a marked stress reaction, (b) by assessing a broad range of physiological indicators, and (c) by considering multiple personality traits as potential moderators. Specifically, we used the Maastricht Acute Stress Test (MAST) to induce acute psychological and physiological stress in 149 healthy adults, randomly assigned to a stress group and a placebo group[26]. Subjectively perceived stress (psychological stress response) was assessed with self-ratings, while heart rate, blood pressure, alpha-amylase and cortisol were obtained as indicators of the physiological stress response. The latter allows for conclusions about the reactivity of both SAM and the HPA. The Big Five personality factors as well as trait anxiety were tested as potential moderators, while the role of general intelligence as a more cognitive but also health-related dimension of personality[27–29], was explored without a concrete hypothesis.

## 2. Methods

### 2.1. Preregistration

Before data analysis, hypotheses, statistical procedures, the sample size, and all variables of interest including their exact operationalization (e.g., which time point was considered) were preregistered in the Open Science Framework: https://osf.io/zd3jf. Note that the present study is part of a larger research project (FOPSID, Fear of Pain, Stress, and Individual Differences in Affect and Cognition) that can be accessed under: https://osf.io/zh3ak/. Note further, that only the first two hypotheses of our preregistration were investigated in the current study (H1, H2), while H3 and H4 will be considered in a following publication.

### 2.2. Hypotheses

**H1:** Based on previous research on the between-subject psycho-physiological correspondence of acute stress markers in laboratory[13] and real-life settings[17] we expect corresponding psychological and physiological indicators of acute stress to show zero or weak positive correlations. Corresponding indicators refer to the matching point in time between psychological and physiological stress indicators, which is (a) at the same time point for blood pressure, heart rate, and alpha-amylase capturing the relatively fast SAM response, and (b) with a 15-20 minute time-delay for cortisol depicting the slower HPA response. Following an existent review[13], two different kinds of correspondence scores are considered: (a) the correspondence between the absolute values of psychological and physiological stress, and (b) the correspondence between the absolute value of psychological stress and the change value in physiological stress, while change values refer to the difference between the absolute value and the average between the first two measurements (before stress onset).

**H2**. We expect the following personality traits to moderate the between-subject psycho-physiological correspondence in acute stress indicators:

H2a. Trait anxiety: Following existing results[14], we expect lower trait anxiety to be related to higher psycho-physiological correspondence.

H2b. Neuroticism: Due to the marked overlap between neuroticism and trait anxiety[30], we initially (see preregistration) expected that lower trait neuroticism is related to higher psycho-physiological correspondence. However, in most recent work of our group, we observed higher neuroticism to be related to higher psychophysiological correspondence[17]. Thus, and in deviation to our preregistration, the association with neuroticism will be tested rather exploratorily.

H2c. Openness: Based on previous findings that body awareness is positively associated with psycho-physiological correspondence[16] and that openness is positively related to body awareness[22], we expect higher openness to be related to higher psycho-physiological correspondence.

H2d. Conscientiousness: Based on previous findings that body awareness is positively associated with psycho-physiological correspondence[16] and that conscientiousness positively relates to body awareness[22], we expect higher conscientiousness to be related to higher psycho-physiological correspondence.

H2e. Extraversion: Based on previous findings[17] and the rationale that extraversion is associated with an orientation towards the outside world[23], we further expect a lower psycho-physiological correspondence for persons with high extraversion scores. Note however, that this hypothesis was not included in our preregistration.

Finally, and as listed in the preregistration under optional analyses, agreeableness and intelligence were also explored as potential moderators. In all analyses, we again differentiate between psycho-physiological correspondence of (a) absolute values and (b) change values.

### 2.3. Participants and sample size justification

The current study is based on data from *N* = 149 participants that were recruited as part of the FOPSID project at Würzburg University. The sample size was determined by a combination of a priori power calculations and what was feasible during the COVID-19 pandemic. Considering prior research, we expected small to medium effect sizes for the main effects hypothesized in the context of the FOPSID project (e.g., correlative relationship between fear of pain and stress *r* = .25[31]). With an alpha level of .05 and a power of .95, this resulted in a sample size of 168 participants (G*Power 3.1[32]). In order to balance the study groups, a sample size of *N* = 180 was initially planned. However, due to restrictions induced by the COVID-19 pandemic, only complete data from 149 participants could be obtained, thus, slightly diminishing the power to .93. Of these 149 participants, 74 persons (mean age = 24.51 years, *SD* = 3.73) were randomly assigned to the stress group, while 75 persons (mean age = 25.04 years, SD = 4.13) were randomly assigned to the control group. The stress group underwent the Maastricht Acute Stress Test (MAST)[26], while the control group participated in a placebo version of the task (Placebo-MAST), where no stress was induced. Note, that we refer to the latter as “mild stress condition” instead of “no stress condition”, as we expect the situation of participating in a psychological experiment not to be completely free of any stressors.

Prior to participation, the following exclusion criteria were assessed via telephone interviews: Age < 18 or > 35; left-handedness; BMI > 30 or < 18; insufficient German language skills (to ensure that instructions are understood); vision and hearing disabilities; Psychology student (beyond the 2nd semester); regular intake of medication affecting the central nervous system; frequent smoking behaviour (> 15 cigarettes per day); drug consumption (including cannabis); history of or acute diagnosis of chronic pain (e.g., headache); episode of severe pain leading to medical treatment within the last six month (e.g., injury); diagnosis of attention-deficit/hyperactivity disorder (ADHD), dyscalculia or dyslexia; acute suffering from any psychological, neurological, cardiovascular, respiratory or endocrinological diseases; disposition for vertigo or fainting. Note that the majority of these criteria were introduced by the fact that the current study belongs to the larger FOPSID project, which includes the assessment of multiple physiological (e.g., cortisol) and psychological (e.g., attentional control) variables, as well as electroencephalographical recordings (EEG) that can be affected by these factors in an uncontrollable way (e.g., cortisol levels can be influenced by smoking, drugs, and medications; attentional control by ADHD diagnosis; EEG by mixed handedness). Furthermore, participants with any history of chronic pain were excluded to prevent any re-activation and harm to a potentially hypersensitive nociceptive system. Participants were instructed to abstain from coffee two hours prior to measurement, to not drink alcohol for 12h before testing, and to immediately inform the study coordinator if any pain medication was taken on the day of testing (either to postpone the measurement or to exclude the participant when medication was to become regular). Experimental procedures were performed in accordance with the declaration of Helsinki, approved by the Institutional Review Board (Psychological Institute, University of Würzburg, Germany, GZEK 2020-18) and informed written consent according to the declaration of Helsinki and to the guidelines of the German Psychological Society was obtained from all participants. Participation in the complete FOPSID project was reimbursed with 10€ per hour, except the Virtual Reality (VR) conditioning experiment (not considered in the current study) that was reimbursed with a fixed amount of money (17.50€) to prevent unintentionally reinforcing avoidance behavior[33].

### 2.4. Study procedure

Figure 1 illustrates the general procedure of the FOPSID project including all measurement time points and the assessed variables. The project consisted of three measurement days (with varying time intervals between days), which were passed by all participants in the same order. On day 1, participants were informed about all study details, informed consent was obtained, all traits were assessed with questionnaires, and intelligence testing took place. On day 2, all participants underwent the Maastricht Acute Stress Test (MAST), a stress induction procedure (or the placebo version of it), before entering an experimental fear conditioning paradigm in VR that was recently developed to investigate the effects of stress on the acquisition and extinction of fear of pain (FoP)[33] in accordance with the Fear-Avoidance Model of chronic musculoskeletal pain[34,35].

**Figure 1.**
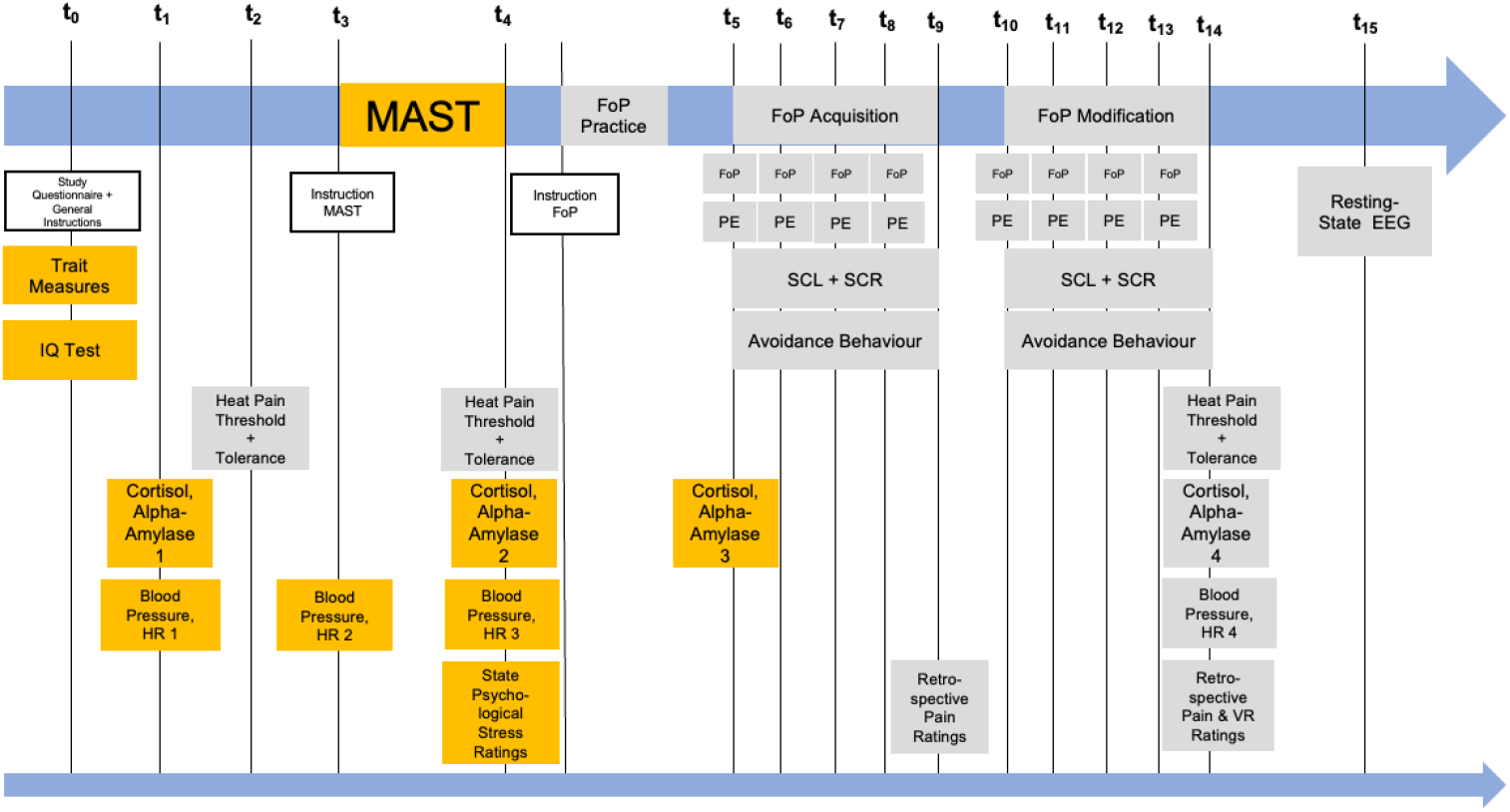
Flowchart of study procedure and assessed variables. Variables analyzed and presented in the current study are highlighted by yellow boxes, while variables not considered here are depicted in grey. Day 1 (t_0_): Study information, personality trait assessments, and intelligence testing. Day 2 (t_1_-t_14_): Stress induction procedure and experimental fear conditioning paradigm in VR. Cortisol and alpha-amylase were assessed four times: In the beginning of the whole experiment (baseline, t_1_), directly after the Maastricht Acute Stress Test (MAST, t_4_), before the acquisition phase of the VR paradigm (exactly 10 min after finishing the MAST, t_5_), and at the end of the experiment (t_14_). Blood pressure and heart rate were also obtained four times: At the beginning (baseline, t_1_), directly before the MAST (t_3_), directly after the MAST (t_4_), and at the end of the experiment (t_14_). Subjective ratings of stress were assessed directly after finishing the MAST (t_4_). Day 3 (t_15_): Electroencephalographical (EEG) resting-state recordings (data not included in the present study). FoP, fear of pain; HR, heart rate; PE, pain expectancy; SCL, skin conductance level; SCR, skin conductance response.

There is general information about e.g. the exact timing of the MAST[26], as well as information about the exact procedure and manipulation checks of the VR paradigm[33]. In brief, the MAST combines physiological and social-evaluative stress exposure by including changing sequences of instructing participants to put their hand in 4°C cold water and counting backwards while receiving negative feedback from the experimenter. After this procedure, the VR fear conditioning paradigm followed for all participants, including a practice phase, a FoP acquisition phase, and a FoP modification or extinction phase. Finally, on day 3, all participants underwent a resting-state EEG recording (data not considered in the present study). While the assessment on day 1 took between 1.5 and 2 hours, the complete experimental procedure on day 2 required 2.5-4 hours (depending on individuals’ choices of movements etc.) and the lab visit for EEG measurements lasted between 1,5 and 2 hours. The psychological and physiological state stress measures obtained during the experimental procedure on day 2 together with the trait measures assessed on day 1 present the input for all analyses of the current study.

### 2.5. Outcome variables

#### 2.5.1. Psychological state stress

Subjectively perceived stress and negative affect were assessed directly after the MAST (reflecting induced stress) or the Placebo-MAST (reflecting mild stress levels without explicit experimental manipulation) (t_4_), respectively. As recommended for the MAST[31], three general items on stress, pain, and unpleasantness were used together with 10 items from the negative affect subscale of the Positive and Negative Affect Schedule (PANAS)[36–38]. The three general items were “How stressful was this procedure right now?” (for stress), “How painful was this procedure right now?” (for pain), and “How unpleasant was this procedure right now?” (for unpleasantness). These items were responded on visual analogue scales, ranging from 0 (*not at all*) to 100 (*extremely*). For the PANAS items, participants were asked to indicate to which extent they felt the different emotions in the current moment, ranging from 1 (*not at all*) to 5 (*very strong*), for example “distressed”, “upset”, or “irritable”. All 13 items combined showed high reliability in our sample (ω = .86 and ω = .85 for the MAST and Placebo-MAST groups, respectively), such that we derived a single state psychological stress score (for details, see the section *Statistical Analyses*).

#### 2.5.2. Physiological state stress

The state stress response of the SAM system was assessed via systolic and diastolic blood pressure, heart rate, and salivary alpha-amylase, while the response of the HPA was examined via salivary cortisol. For most comprehensive insights and as outlined in our preregistration in detail, all variables were operationalized in two ways: a) By absolute scores obtained when the stress response is assumed to be maximal and b) by change scores calculated as difference between this score and a baseline measurement.

##### Heart rate

Heart rate (HR) was assessed in beats per minute using the Sanitas SBM 21 (Breuer GmbH, Ulm, Germany) oscillometric blood pressure monitor with arm cuff. Heart rate values obtained at the end of the stress induction procedure, i.e., at the same time point as psychological stress ratings (t_4_), were considered as absolute scores (HR_abs_), while the difference between this score and a baseline score (average across both time points before the stress induction procedure: t_1_, t_3_) served as change score (HR_change_).

##### Blood pressure

Systolic and diastolic blood pressure (BP) in mmHg were also measured using the Sanitas SBM 21 (Breuer GmbH, Ulm, Germany) oscillometric BP monitor. Similarly as for heart rate, diastolic and systolic blood pressure obtained at the end of the stress induction procedure (t_4_) served as absolute scores (BP_abs_), while change scores (BP_change_) were calculated as differences between these scores and the baseline (again averaged across both time points before the stress induction procedure: t_1_, t_3_).

##### Alpha-amylase

The same definitions and timing of absolute (AA_abs_) and change scores (AA_change_ = t_4_ – (t_1+_t_3_)/2) refer also to salivary alpha-amylase.

##### Cortisol

Due to the slower HPA response to stress, cortisol values assessed exactly 10 minutes after finishing the stress induction (t_5_) served as absolute scores. Change scores were calculated as the difference between this score and the baseline at t_1_.

Concentrations of alpha-amylase and cortisol were assessed by salivary samples and analyzed in an external laboratory at Dresden University. It was ensured that all saliva samples kept frozen at -20°C until analysis. Concentrations of alpha-amylase in saliva were measured by an enzyme kinetic method: Saliva was processed on a Genesis RSP8/150 liquid handling system (Tecan, Crailsheim, Germany). First, saliva was diluted 1:625 with double-distilled water by the liquid handling system. Twenty microliters of diluted saliva and standard were then transferred into standard transparent 96-well microplates (Roth, Karlsruhe, Germany). Standard was prepared from ‘‘Calibrator f.a.s.’’ solution (Roche Diagnostics, Mannheim, Germany) with concentrations of 326, 163, 81.5, 40.75, 20.38, 10.19, and 5.01 U/l alpha-amylase, respectively, and bidest water as zero standard. Afterwards, 80 ml of substrate reagent (*a*-amylase EPS Sys; Roche Diagnostics, Mannheim, Germany) was pipetted into each well by using a multichannel pipette. The microplate containing sample and substrate was then warmed to 37°C by incubation into a waterbath for 90 s. Immediately afterwards, a first interference measurement was obtained at a wavelength of 405 nm using a standard ELISA reader (Anthos Labtech HT2, Anthos, Krefeld, Germany). The plate was then incubated for another 5 min at 37°C in the waterbath, before a second measurement at 405 nm was taken. Increases in absorbance were calculated for unknowns and standards. Increases of absorbance of diluted samples were transformed to alpha-amylase concentrations using a linear regression calculated for each microplate (Graphpad Prism 4.0c for MacOSX, Graphpad Software, San Diego, CA).

For the analyses of cortisol concentrations, salivettes were first centrifuged at 3,000 rpm for 5 min after thawing, which resulted in a clear supernatant of low viscosity. The exact concentrations were then derived by using commercially available chemiluminescence immunoassay with high sensitivity (IBL International, Hamburg, Germany). All intra- and interassay coefficients for alpha-amylase and cortisol were below 9% indicating good measurement quality. More details about the analytical procedure can be seen elsewhere[39].

### 2.6. Moderator variables

#### 2.6.1. Trait anxiety

Trait anxiety was assessed together with all other trait variables prior to the experimental manipulation on a separate day (t_0_) using the trait anxiety subscale (α = .88) from the German version of the State-Trait-Depression-Inventory (STADI)[40]. The trait part of the STADI includes 20 items consisting of statements about the person, e.g., “I enjoy life”. Participants were asked to indicate on a 4-point scale how often each statement holds true from 1 = “not at all” to 4 = “a lot”.

#### 2.6.2. Personality traits

The Big Five personality traits (openness, conscientiousness, extraversion, agreeableness, neuroticism) were also assessed at t_0_ using the German version[41]of the NEO-FFI[42]. The questionnaire consisted of 60 items, measuring each trait with 12 items. These items were statements about how the participants perceive themselves, e.g., “I am not easily disturbed”, and were rated on 5-point Likert scales ranging from 0 (*strongly disagree*) to 4 (*strongly agree*). Scale reliabilities were reported to range between α = .72 and α = .87[41].

#### 2.6.3. Intelligence

Finally, also general intelligence was obtained at t_0_ with the German version of Raven’s Advanced Matrices (APM)[43,44]. The APM is a non-verbal intelligence assessment intended to capture general intelligence relatively independent of culture, knowledge and previous education. Note that the advanced rather than the standard version was used to best differentiate also in the higher ability range as can be expected for a primarily student sample. For all further analyses, the total number of correct responses was used (APM sum score). Reliability was reported to be α = .85[43].

### 2.7. Statistical analyses

All analyses were performed using the software R[45]. Specifically, linear and moderated regressions were implemented with the lm() function and 95% confidence intervals were obtained with the confint() function. In doing so, we determine psychophysiological correspondence on the between-subject level, i.e., the correspondence of psychological and physiological measures between different persons, so that persons scoring high on psychological measures tend to score also high on physiological measures. Prior to analyses, outliers (values deviating more than three standard deviations from the sample mean) were replaced with values two standard deviations from the sample mean (this step was taken in accordance with an analogous procedure[31]). Due to largely differing value ranges across psychological and physiological variables, all scores were tested for normal distribution using the Shapiro-Wilk test as well as graphical inspections. As almost all distributions were non-normal, Spearman’s ρ was used as correlation coefficient. Further, all scores were *z*-standardized before entering the multiple regression models, while Table 1 illustrates the unstandardized scores.

**Table 1.** Descriptive statistics and difference tests for MAST and Placebo-MAST groups.

|  | MAST |  |  |  | Placebo-MAST |  |  |  | Welch's *t*-test |  |  |  |
| --- | --- | --- | --- | --- | --- | --- | --- | --- | --- | --- | --- | --- |
| | *M* | *SD* | Min, Max | $\omega$ | *M* | *SD* | Min, Max | $\omega$ | *t* | df | *p* | *d* |
| Age | 24.51 | 3.73 | 18, 35 | -- | 25.04 | 4.13 | 18, 34 | -- | 0.82 | 145.82 | .416 | 0.13 |
| **Psychological variables** |  |  |  |  |  |  |  |  |  |  |  |  |
| State psychological stress<sup>m</sup> | 0.61 | 0.83 | -0.97, 2.86 | -- | -0.60 | 0.57 | -1.19, 1.88 | -- | -10.29** | 129.04 | <.001 | 1.69 |
| Trait-Anxiety | 2.13 | 0.30 | 1.30, 2.80 | .91 | 2.12 | 0.28 | 1.65, 2.94 | .92 | -0.13 | 145.57 | .899 | 0.02 |
| Neuroticism | 1.44 | 0.63 | 0.25, 3.17 | .87 | 1.55 | 0.64 | 0.08, 2.92 | .88 | 1.06 | 147 | .292 | 0.17 |
| Openness | 2.73 | 0.47 | 1.33, 3.75 | .75 | 2.74 | 0.50 | 1.50, 3.83 | .78 | 0.12 | 146.68 | .908 | 0.02 |
| Conscientiousness | 2.56 | 0.71 | 0.83, 4.00 | .91 | 2.59 | 0.63 | 1.08, 3.83 | .90 | 0.26 | 144.55 | .796 | 0.04 |
| Extraversion | 2.33 | 0.58 | 1.08, 3.75 | .86 | 2.46 | 0.61 | 0.92, 3.58 | .88 | 1.29 | 146.75 | .201 | 0.21 |
| Agreeableness | 2.56 | 0.56 | 1.10, 3.67 | .81 | 2.65 | 0.55 | 1.36, 4.00 | .80 | 1.03 | 146.86 | .304 | 0.17 |
| Intelligence | 27.54 | 4.66 | 14.00, 35.00 | -- | 26.29 | 4.27 | 17.00, 36.00 | -- | -1.70 | 145.48 | .091 | 0.28 |
| **Physiological variables** |  |  |  |  |  |  |  |  |  |  |  |  |
| HR<sub>change</sub><sup>m</sup> | -5.65 | 10.18 | -41.36, 17.00 | -- | 0.70 | 14.00 | -50.00, 64.50 | -- | 3.17 | 135.19 | .999 | 0.52 |
| HR<sub>abs</sub><sup>m</sup> | 76.00 | 14.35 | 46.00, 120.95 | -- | 80.68 | 18.05 | 51.00, 123.00 | -- | 1.75 | 140.64 | .959 | 0.29 |
| BP<sub>change</sub><sup>m</sup> | 4.21 | 6.71 | -10.27, 19.69 | -- | -1.96 | 7.16 | -23.84, 20.50 | -- | -5.42** | 146.63 | <.001 | 0.89 |
| BP<sub>abs</sub><sup>m</sup> | 106.97 | 9.25 | 81.83, 124.95 | -- | 100.47 | 8.82 | 74.65, 120.89 | -- | -4.39** | 146.47 | <.001 | 0.72 |
| AA<sub>change</sub><sup>m</sup> | 31.80 | 50.60 | -56.80, 157.73 | -- | -3.19 | 43.45 | -128.11, 134.28 | -- | -4.48** | 140.78 | <.001 | 0.74 |
| AA<sub>abs</sub><sup>m</sup> | 128.26 | 107.77 | 7.15, 472.16 | -- | 92.55 | 94.65 | 4.42, 453.85 | -- | -2.13* | 141.64 | .018 | 0.35 |
| Cortisol<sub>change</sub><sup>m</sup> | 5.64 | 6.43 | -9.79, 23.67 | -- | -1.44 | 3.08 | -10.94, 4.73 | -- | -8.55** | 104.49 | <.001 | 1.40 |
| Cortisol<sub>abs</sub><sup>m</sup> | 10.88 | 6.41 | 1.27, 25.49 | -- | 4.20 | 2.62 | 0.48, 13.76 | -- | -8.32** | 96.56 | <.001 | 1.37 |

*Note.* <sup>m</sup> = psychological and physiological variables that capture the stress reaction following the experimental manipulation in the MAST group and are thus assumed to differ between groups (i.e., manipulation check). Statistical significance of group differences was assessed with Welch's *t*-tests. For variables assumed to differ between groups, Welch's *t*-tests were conducted one-sided, while all other variables were tested two-sided. Change scores (indicated by subscript) were calculated such that the baseline score was subtracted from the score following stress induction or Placebo, respectively (i.e., score after experimental manipulation – baseline score), that is, a positive value indicates an increase in stress following experimental manipulation, while a negative value indicates a decrease in stress. Note that while this table presents values in their original measurement scale, all variables were z-standardized before entering statistical regression models, testing our main hypotheses. Sample size (*N*) of the MAST group = 74; *N* of the Placebo-MAST group = 75. \* = significant at $p < .05$ , \*\* = significant at $p < .001$ (uncorrected for multiple comparisons). HR = heart rate; BP = blood pressure; AA = alpha amylase.

To obtain a composite score for state psychological stress, the ten negative affect items of the PANAS and the three single items assessing stress, pain, and unpleasantness (see *Measures* for details) were combined into a single composite score of psychological stress (PS) after running an exploratory factor analysis using the fa() function. To obtain a single score for blood pressure, we calculated the mean arterial pressure based on systolic and diastolic blood pressure using a provided formula[46]. In a post-hoc analysis, which was not pre-registered, we obtained a composite score for state physiological stress analogously to the composite score for state psychological stress, i.e., by obtaining the factor score of an exploratory factor analysis based on the four physiological variables. All analyses were rerun with that latent physiological stress factor as variable of interest to gain insight into a potential superiority of a composite physiological score as compared to separate physiological stress indicators. The MAST and Placebo-MAST groups were analyzed separately. Respective models are presented in the results section. Statistical significance was assumed at *p* < .05. Furthermore, effect sizes and their 95% confidence intervals were computed. To account for multiple comparisons, all *p*-values were corrected with the Benjamini-Hochberg correction (α = .05) within the same model across the MAST and Placebo-MAST conditions, respectively[47]. Since the Holm-Bonferroni-Method has been evaluated as too conservative, especially when it comes to exploratory analyses, we considered the Benjamini-Hochberg correction as the most appropriate method (instead of the Holm-Bonferroni[48], which was initially specified in our preregistration). For most comprehensive insights and to enhance clarity, both corrected and uncorrected *p*-values are reported and the exact model specifications are described in the beginning of the respective results sections.

Bayesian regression analyses, which was not preregistered, but are more suitable to quantify evidence for the null hypothesis (i.e., evidence against a correspondence between psychological and physiological stress indicators) than the frequentist framework of significance testing [49–51] was implemented in JASP[52]. All Bayesian analyses were controlled for age and employed Jeffreys–Zellner–Siow priors with the default Cauchy scale (*r* = .354).

### 2.8. Data and code availability

All analysis code can be accessed at the Open Science Framework under: https://osf.io/zh3ak/files/osfstorage. The data can be obtained from the authors by request.

## 3. Results

### 3.1. Descriptive statistics and manipulation checks

Descriptive statistics and reliabilities for each variable and each group (MAST vs. Placebo-MAST) are listed in Table 1. As indicated by McDonald’s ω coefficients, internal consistencies ranged between ω = .75 and ω = .92 and were remarkably similar across the MAST and the Placebo-MAST group.

To examine whether the stress induction procedure was successful one-sided Welch’s *t*-tests were conducted for psychological and physiological state stress measures, comparing the MAST group with the Placebo-MAST group. For variables for which no group differences were assumed (e.g., personality traits), two-sided Welch’s *t*-tests were performed. As depicted in Table 1, all psychological and physiological state stress measures differed significantly between the two groups in the expected direction (higher stress in the MAST group) with mostly moderate to large effect sizes (uncorrected for multiple comparisons[53]). The only exception were heart rate values, that did not differ significantly between both groups. In sum, these manipulation checks suggest successful stress induction.

Spearman correlations among all study variables for the MAST and the Placebo-MAST group, are listed in Supplementary Tables S1 and S2. As expected, psychological and physiological stress indicators were not significantly correlated with each other. Neuroticism was the only Big Five personality trait which significantly correlated with state psychological stress in both experimental groups (ρ = .26, *p* = .028 and ρ = .34, *p* = .003 in the MAST and the Placebo-MAST group, respectively; uncorrected), while trait anxiety was only significantly correlated with state psychological stress in the Placebo-MAST group (ρ = .27, *p* = .020) and all other personality traits demonstrated no significant association. Nearly all physiological stress indicators were significantly positively inter-correlated).

### 3.2. Psycho-physiological correspondence (H1)

To test our first hypothesis that psychological and physiological stress indicators show zero or only weak positive correlations, we first conducted linear regression analyses with the latent state psychological stress factor as outcome variable, and one physiological variable and age as predictor variables (Models 1-8 in Supplementary Table S3). Out of all physiological variables, only cortisol demonstrated a statistically significant relationship with psychological stress, while this applies only for the MAST group and only when considering uncorrected *p*-values (Model 4; β = 21, *p* = .024). Accordingly, Model 4 (MAST) showed the highest *R*^*2*^ of .11 which was statistically significant with *F*(2,71) = 4.38; *p* = .016. All other models were not significant with a mean *R*^*2*^ of .04. Secondly, we ran two comprehensive models for each group, comprising all physiological (absolute or change, respectively) variables and age as predictors, and psychological stress as outcome (see Table 2). Absolute and change values of cortisol were the only significant predictors of psychological stress besides the covariate age, but again only in the MAST group and only when considering uncorrected *p*-values (Model 9; β = .24, *p* = .016, *R*^*2*^ *=* .14; Model 10; β = .21, *p* = .041, *R*^*2*^ *=* .11). In sum, we found no significant correspondence between psychological and physiological stress indicators irrespective of whether stress was induced or not and irrespective of the physiological stress indicator considered with the slight exception of cortisol.

**Table 2.** Multiple regression models on psycho-physiological correspondence (H1)

|  | State psychological stress (MAST) |  |  |  |  | State psychological stress (placebo-MAST) |  |  |  |  |
| --- | --- | --- | --- | --- | --- | --- | --- | --- | --- | --- |
| | $\beta$ (SE) | *p* | *p*<sub>adj</sub> | 95% CI | *R*<sup>2</sup> (*p*) | $\beta$ (SE) | *p* | *p*<sub>adj</sub> | 95% CI | *R*<sup>2</sup> (*p*) |
| **Model 9** |  |  |  |  | .14<br>(.067) |  |  |  |  | .08<br>(.357) |
| HR<sub>abs</sub> | .15 (0.11) | .169 | .339 | [-.07, .37] |  | .10 (0.06) | .102 | .246 | [-.02, .23] |  |
| BP<sub>abs</sub> | .00 (0.10) | .963 | .963 | [-.21, .20] |  | .03 (0.07) | .649 | .779 | [-.11, .18] |  |
| AA<sub>abs</sub> | -.11 (0.10) | .273 | .413 | [-.30, .09] |  | -.08 (0.08) | .275 | .413 | [-.23, .07] |  |
| Cortisol<sub>abs</sub> | .24 (0.10) | .016* | .065 | [.05, .44] |  | .05 (0.16) | .780 | .851 | [-.28, .37] |  |
| Age | -.22 (0.10) | .039* | .117 | [-.43, -.01] |  | -.06 (0.07) | .393 | .524 | [-.20, .08] |  |
| **Model 10** |  |  |  |  | .11<br>(.149) |  |  |  |  | .06<br>(.556) |
| HR<sub>change</sub> | .05 (0.12) | .702 | .766 | [-.20, .29] |  | -.01 (0.06) | .882 | .882 | [-.13, .11] |  |
| BP<sub>change</sub> | -.13 (0.11) | .271 | .464 | [-.35, .10] |  | .03 (0.07) | .693 | .766 | [-.12, .17] |  |
| AA<sub>change</sub> | -.09 (0.10) | .364 | .515 | [-.29, .11] |  | -.07 (0.08) | .386 | .515 | [-.23, .09] |  |
| Cortisol<sub>change</sub> | .21 (0.10) | .041* | .122 | [.01, .41] |  | -.16 (0.14) | .257 | .464 | [-.44, .12] |  |
| Age | -.24 (0.11) | .032* | .122 | [-.45, -.02] |  | -.08 (0.07) | .228 | .464 | [-.21, .05] |  |

*Note.* Absolute scores (indicated by the subscript abs) represent stress scores after experimental stress induction (MAST) or time-corresponding values in the control group, respectively (Placebo-MAST). Change scores (indicated by subscript) were calculated such that the baseline score was subtracted from the score following stress induction or Placebo, respectively (i.e., score after experimental manipulation – baseline score). \* = significant at $p < .05$ , \*\* = significant at $p < .001$ . HR = heart rate; BP = blood pressure; AA = alpha amylase; $\beta$ = standardized regression coefficient; SE = standard error; *p*<sub>adj</sub> = Benjamini-Hochberg corrected *p*-values.

While these findings are consistent with our first hypothesis, the absence of significant effects does not constitute definitive evidence for the null hypothesis. Therefore, additional not preregistered Bayesian regression analyses were conducted. In the MAST group (see Table 3), Bayes factors comparing the null model to models including one of the four physiological indicators and age yielded BF□ □ values between 1.00 and 3.31, thus providing weak to moderate evidence in favor of the null hypothesis [54] i.e., absence of psycho-physiological correspondence. In the Placebo-MAST group, Bayes factors were consistently larger, with BF□ □ values ranging from 3.40 to 7.13, thus providing moderate evidence for the null hypothesis. Consistent with the frequentist analyses, explained variance (R^2^) was low across all models (R^2^ ≤ .04), and no physiological variable or age emerged as reliable predictor of psychological stress. These results provide quantitative support for the absence of a psycho-physiological correspondence.

**Table 3.**
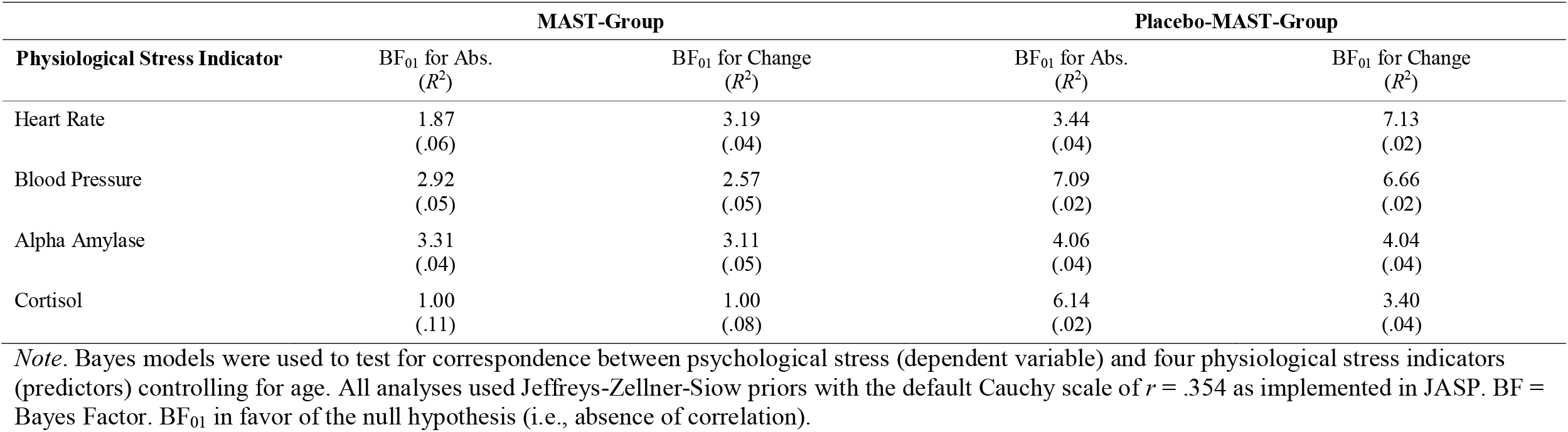
Bayes regression analyses on psycho-physiological correspondence (H1)

### 3.3 Moderators of the psycho-physiological correspondence (H2)

To evaluate our second hypothesis (personality traits moderate the psycho-physiological correspondence), moderated regression analyses were conducted, including all absolute or change indicators of physiological stress, respectively, the covariate age, the respective trait of interest (anxiety, openness, conscientiousness, neuroticism, extraversion, agreeableness, and intelligence) as well as the interaction terms between the respective trait of interest and all physiological variables (Models 11-24). To enhance readability across the 14 models, Table S4 includes only the moderator effects, while Table S5 contains the corresponding main effects of the physiological variables as well as age. For explorative purposes, effects whose uncorrected *p*-value reached statistical significance are also reported in the following text together with effects that reached significance when correcting for multiple comparison (*p*_*adj*_).

Our first sub-hypothesis positing that lower trait anxiety relates to higher psycho-physiological correspondence (H2a) applied for the correspondence between psychological stress and blood pressure in the MAST group (Model 11; β = -.24, *p* = .014) as well as for the correspondence between psychological stress and alpha-amylase in the Placebo-MAST group (Model 11; β = -.16, *p* = .014). However, in the MAST group, we also identified a moderator effect of trait anxiety in the opposite direction as for alpha-amylase (Model 11; β = .17, *p* = .038). Overall, the *R*^*2*^ of this Modell was .26 and the model was statistically significant with *F*(10,62) = 2.16; *p* = .033. No significant moderator effects for trait anxiety were observed in respect to the other physiological stress indicators (all *p* > .05; Table S4).

Second, we explored the relation between neuroticism and psycho-physiological correspondence as previous evidence was contradictory (H2b). Model 17 indicated a positive relation in the Placebo-MAST group only, i.e., higher neuroticism was associated with higher correspondence between psychological stress and absolute heart rate (Model 17; β = .15, *p* = .031).

Third, we expected higher openness to be related to higher psycho-physiological correspondence (H2c). In line with this hypothesis, higher openness was positively associated with higher correspondence between psychological stress and absolute alpha-amylase (Model 13; β = .37, *p* = .002, *p*_*adj*_ = .011) as well as to higher correspondence between psychological stress and the change in alpha-amylase (Model 14; β = .34, *p* = .009). Both such moderation effects apply to the MAST group only. Contrary to our hypothesis, higher openness was associated with lower correspondence between psychological stress and absolute cortisol in the MAST group (Model 13; β = -.26, *p* = .009), while no significant moderation effects were observed for the Placebo-MAST group and in respect to the other physiological indicators (all *p* > .05; Table S4). Overall, Model 13 (MAST) reached statistical significance with *R*^*2*^ = .31 (*F*(10,62) = 2.74; *p* = .007).

Fourth, we proposed higher conscientiousness to be related to higher psycho-physiological correspondence (H2d). Supporting this hypothesis, conscientiousness was associated with higher correspondence between psychological stress and absolute blood pressure (Model 15; β = .23, *p* = .021) as well as to a higher correspondence between psychological stress and the change in cortisol (Model 16; β = .29, *p* = .003, *p*_*adj*_ = .017). Again, both were true for the MAST group only, i.e., after stress was induced. Contrary to our expectations, conscientiousness was associated with lower correspondence between psychological stress and the change in alpha-amylase in the MAST group (Model 16; β = -.32, *p* = .002, *p*_*adj*_ = .015). Overall Model 16 (MAST) reached a *R*^*2*^ of .31 which was statistically significant with *F*(10,62) = 2.8 (*p* = .006).

Finally, we explored extraversion, agreeableness and intelligence as potential moderators. For extraversion and intelligence, not any moderation effect reached statistical significance (see Models 19, 20, 23, and 24 in Table S4). For agreeableness, only one association was statistically significant: Higher agreeableness was related to a higher correspondence between psychological stress and absolute heart rate in the MAST group (Model 21; β = .29, *p* = .011).

### 3.4. Post-hoc analyses: A composite score of physiological stress

To investigate whether a composite score of physiological stress might yield additional insights into psycho-physiological correspondence, we repeated all analyses using aggregated absolute and change physiological stress scores, respectively (not preregistered). Descriptive statistics are listed in Table S6, and correlations to all other study variables are displayed in Table S7. Notably and in clear contrast to the psychological stress variables, the physiological stress variables showed unsatisfying reliability coefficients (ω = .12 to ω = .45), suggesting low internal consistency of the four indicators heart rate, blood pressure, alpha-amylase and cortisol irrespective of whether stress was induced or not. Moderation effects between traits of interest and the psycho-physiological correspondence considering this composite score of physiological stress are presented in Table S8 (Models 25 to 40). Across all models, no single moderation effect reached statistical significance.

## 4. Discussion

Although a correspondence between psychological and physiological stress indicators seems evolutionary advantageous[12] and was proposed as relevant for physical and mental health[14], research suggests rather low or non-existent correspondence between psychological and physiological indicators of acute stress[13,55]. The current study provides a comprehensive test of this correspondence. It closes the gaps of previous research (a) by applying a stress induction procedure that comprises exposure to rather high psychological *and* physiological stressors as well as a control condition in which no stress was induced and (b) by considering four physiological indicators of stress (blood pressure, heart rate, alpha-amylase, cortisol) as well as multiple personality traits (trait anxiety, the Big Five, intelligence) as potential moderators.

In line with our first hypothesis, we did not observe any significant correspondence between psychological and physiological stress indicators after controlling for multiple testing and Bayesian analyses provided quantitative evidence for the null hypothesis, i.e., absence of any psycho-physiological correspondence. When considering uncorrected significance, cortisol was the sole physiological stress indicator correlated with psychological stress - in the MAST group only. This is at least partially in accordance with a meta-analysis which observed significant correlations between cortisol and psychological stress indicators in 25% of the studies involving the TSST[13]. At least in respect to our study, one may speculate, however that the potential cortisol-specificity of the correspondence may result from a closer timing of both assessments, as psychological stress was measured directly after the MAST (10 min after the acute stress induction started) together with the expected maximum of cortisol[26]. Further, cortisol release is modulated by the activity of the HPA axis, which contrasts with the other obtained physiological stress indicators reflecting SAM activity. However, as the observed association did not pass the corrected significance threshold and Bayes factors still favor the null hypothesis, further research including larger samples is essentially required to elucidate the potential cortisol-specificity (or HPA axis-specificity) of the correspondence. Finally, the uncorrected cortisol-specific effect refers to models testing for psycho-physiological correspondence in the MAST group only, suggesting the induction of a marked stress response as an important prerequisite for validly studying psycho-physiological correspondence. On the one hand, physiological stress symptoms below a certain threshold might be difficult to notice (e.g., a heart rate increase), and thus can hardly affect self-reported (psychological) stress ratings. On the other hand, psycho-physiological correspondence is evolutionary most functional under high stress, as physiological and psychological stress reactions running in parallel increase the probability to survive actions[56].

Concerning our second hypothesis, proposing individual differences in basic personality traits as moderators of the psycho-physiological correspondence, eleven significant interactions were observed before controlling for multiple testing out of which six were in the expected direction and three passed the corrected *p*-threshold (two in the expected direction): In line with our hypotheses conscientiousness and openness showed four significant interaction effects indicating higher conscientiousness/openness to be related to higher correspondence. Two out of these effects stayed significant even after controlling for multiple testing. These two personality traits, however, also showed two interaction effects in the opposite direction out of which one survived the correction for multiple testing. A further contra-hypotheses interaction effect was identified for neuroticism (higher correspondence in persons scoring higher in neuroticism) while two expected effects (higher correspondence in persons scoring higher trait anxiety) and one effect in the opposite direction emerged for trait anxiety as well as one unexpected effect for agreeableness (higher correspondence in persons with lower agreeableness scores). Of note, all effects regarding neuroticism, trait anxiety and agreeableness did not pass the significance threshold when correcting for multiple testing and no effects were observed for extraversion and general intelligence. Regarding the four investigated physiological stress indicators alpha-amylase and cortisol were involved in the three interaction effects that stayed significant after controlling for multiple testing.

A moderation effect of conscientiousness on the psycho-physiological correspondence has also been observed previously[17] and may be explained by the higher body awareness of conscientious persons[22]. This is consistent with evidence that both conscientiousness[57,58] and psycho-physiological correspondence[14] are predictive of physical and mental well-being. However, we did not expect to observe an opposite effect with respect to alpha-amylase change scores. Potentially, this could be explained by both stress axes (SAM, HPA) differing in their timing and, most importantly, potentially also in which direction higher conscientiousness influences their responsivity. Specifically, it can be speculated that higher conscientiousness facilitates the alignment of psychological stress ratings and HPA reactivity, while it may reduce the alignment between psychological stress ratings and SAM responses for which alpha-amylase might be a more direct marker compared to heart rate and blood pressure for which no effects were observed. However, this very vague interpretation was proposed post-hoc and has to be tested in future studies comprehensively. The significant interaction effect of openness, indicating higher correspondence between psychological stress and absolute alpha-amylase in persons scoring higher on openness was again in the direction expected based on former research demonstrating higher openness to be related to higher body awareness. However, if our above-outlined interpretation of the contradictory effect of alpha-amylase change scores in respect to conscientiousness holds true, this finding would reveal an important distinction between the consideration of change versus absolute scores of alpha-amylase that has to be taken into account in future research.

Notably, nine out of the 11 uncorrected significant effects were observed in the MAST group only, i.e., after exposure to marked psychological (social-evaluative) and physiological (cold water) stress. Thus, it seems that only under a certain amount of stress, personality factors come into play (conscientiousness, openness) and align psychological and physiological stress responses. Personality theories which do *not* conceptualize traits as fully decontextualized constructs provide potential interpretations for this observation (Cybernetic Big Five Theory[59]; Trait Activation Theory[60]). When we consider psycho-physiological correspondence as trait, high stress might present the trait-relevant situation for individual differences in the correspondence (inclusively their moderation) to become visible. Although this has to be further investigated in future studies, we speculate that personality-related individual differences in interoception in general, and in particular in the ability to notice and become aware of bodily stress symptoms might play a critical role.

This study has limitations but provides important insights for future research on the psycho-physiological correspondence. First, only 2 out of 80 hypotheses regarding the potential moderating effects of personality traits on the psycho-physiological correspondence were supported after correcting for multiple testing and 9 out of 80 hypotheses were confirmed when considering uncorrected *p*-values. Thus, most of our hypotheses were actually *not* supported. At the same time, it would be premature to conclude that personality traits do not moderate the correspondence at all. Rather our study showcases that potential moderating effects of personality traits may depend on (a) the respective trait, (b) the measures for psychological and physiological stress, (c) the intensity, duration and type of the stress and (d) the calculation of the psycho-physiological correspondence (within or between subjects). Therefore, we recommend future studies to evaluate these factors systematically to allow for more specific hypotheses in future research. For example, more specific personality traits or trait-like skills such as body awareness or interceptive abilities that are more directly related to the perception of physiological stress symptoms should be included. In addition, the notion of psycho-physiological correspondence implicitly assumes an association between all kinds of psychological and physiological stress markers. However, this assumption is much too undifferentiated both in terms of the psychological and the physiological stress markers. One and the same score on a psychological stress scale could stem from or be associated with different emotions (e.g., anxiety, anger) and it seems therefore reasonable or even mandatory for future studies to operationalize the construct of psycho-physiological correspondence in a much more fine-grained way. This would result in several correspondence types between specific self-reported stress-related emotions and specific physiological stress indicators, each to be considered separately and to be tested for potential moderation effects of personality traits. Since our study was the first one to investigate the psycho-physiological convergence in a stress and in a placebo condition and as we found at least some preliminary evidence that the correspondence was more pronounced under stress, it also seems worthwhile to investigate such different types of psycho-physiological correspondences in different more or less stressful contexts and over shorter versus longer time periods.

Another interesting question for future research is whether the assumption of high evolutionary functionality of psycho-physiological correspondence[12] is valid under all circumstances. For example, rescue workers may experience high levels of physiological stress but may have to suppress the respective signals on the psychological level to complete life-saving emergency medical procedures. Further, the human stress system evolved to handle immediate physical threats in primeval times by preparing the body for “fight or flight.” In contemporary society, most stressors are psychological – such as work pressures, financial concerns, and social dynamics. This shift implies that the physiological stress response is often activated in situations where physical action is neither necessary nor appropriate. For example, you would not run away or physically attack your boss when experiencing an argument with him/her. Therefore, while the multi-modal stress response was advantageous in ancestral environments for dealing with acute threats, its frequent activation by modern, can be maladaptive, suggesting a mismatch between our evolved stress mechanisms and contemporary cultural environments. As a consequence, the psycho-physiological correspondence would have to be complemented by a social component meaning that persons should be able to transform the primeval “fight or flight” reaction in behaviors that are compatible with societal norms which have evolved during the cultural evolution[61].

Our contribution may also stimulate an extension of the well-established Exposure-Reactivity Model of stress and coping[19]. In this model, personality traits are conceptualized as possible moderators between stressor exposure and the stress reaction. Individual variations in the correspondence between different response systems of stress, i.e., the psycho-physiological correspondence addressed in this paper, could also be integrated in this model. Finally, the above suggested avenues for future research should be complemented by intensive longitudinal studies focused on single participants. Such idiographic investigations allow detecting potential idiosyncratic correspondence patterns: For some individuals there may be a correspondence with regard to certain psycho-psychological pairings (e.g., psychological stress and cortisol) but not for other pairings (e.g., psychological stress and heart rate) and there may be individuals for whom there is no correspondence at all. Resulting idiosyncratic correspondence types may then be aggregated to clusters across individuals showing similar patterns and can be investigated further, for instance, with regard to relations with personality traits.

To conclude, we recommend future studies to (a) consider a broad range of both psychological and physiological stress indicators, (b) to investigate specific types of psycho-physiological correspondence resulting from certain pairings of indicators, (c) to include concepts reflecting interoceptive abilities and body awareness, (d) to systematically vary the intensity and duration of stressors, and (e) to combine large-sample nomothetic approaches with longitudinal idiographic research.

## 5. Conclusion

This study is the first to comprehensively investigate the potential moderating role of basic personality traits on the correspondence between acute psychological stress and multiple physiological stress markers. In extension to previous research, we used an experimental paradigm that included both the induction of psychological and physiological stress (MAST) as well as a control condition in which no stress was induced (Placebo-MAST). Although most of the hypothesized moderations failed to reach statistical significance, and some interaction effects turned out to contradict our hypotheses, our study presents promising starting points for future research and reveals that the specific situational circumstances as well as participants’ personality traits need to be taken into account when studying the psycho-physiological correspondence of stress indicators. For future research, we propose (a) the consideration of more specific personality traits or trait-like skills with regard to different corresponding pairings of stress-specific emotions and specific physiological indicators, (b) the systematic variation of stress intensity and laboratory vs. real life contexts, and (c) the combination of nomothetic and idiographic analyses, as promising avenues to disentangle the mystery of the missing psycho-physiological correspondence, as critically impacting mental health and human well-being.

## Acknowledgement

KHR is supported by dtec.bw - Digitalization and Technology Research Center of the Bundeswehr. dtec.bw is funded by the European Union – NextGenerationEU. Data acquisition (and the whole the FOPSID project) was funded by Würzburg University. The Open Access publication of this article is covered by the Open Access Publication Fund of the University of Bundeswehr Munich. The authors thank all participants for their participation, Prof. Paul Pauli for providing facilities and for supporting data acquisition, as well as Amelie Schirmer, Dorna Marzban, Kilian Rolle, Annika Aster, Ines Jetzinger, and Katharina Schmelz for their help with participant management and data acquisition.

## CRediT authorship contribution statement

**Kirsten Hilger:** Conceptualization, Methodology, Formal Analysis, Project Administration, Writing – Original Draft, Supervision, Funding acquisition. **Irma Talic:** Conceptualization, Formal Analyses, Writing – Original Draft. **Karl-Heinz Renner:** Conceptualization, Writing – Original Draft, Supervision, Funding.

## Ethics Approval Statement

All experimental procedures were in accordance with the declaration of Helsinki and approved by the local ethics committee (Psychological Institute, University of Würzburg, Germany, GZEK 2020-18). Informed written consent according to the declaration of Helsinki and to the guidelines of the German Psychological Society was obtained from all participants.

## Supplementary Material

Supplementary Material associated with this article can be found in a separate document.

