## Supplementary for "Individual Differences in the Correspondence Between Psychological and Physiological Stress Indicators"

**Author Note:**

Main Text: Max. 4500 words, now: 6811 – 6361 without Hypotheses.

*** Corresponding Authors:**

Dr. Kirsten Hilger, Department of Psychology I, Marcusstr. 9-11, D-97070 Würzburg

Prof. Dr. Karl-Heinz Renner, Department of Psychology, Werner-Heisenberg-Weg 39 D-85577 Neubiberg,

**Table S1**

*Correlations between all study variables*

| Variable | 1 | 2 | 3 | 4 | 5 | 6 | 7 | 8 | 9 |
|---|---|---|---|---|---|---|---|---|---|
| 1. State psychological stress | --- | | | | | | | | |
| 2. Openness | .00 /.07 | --- | | | | | | | |
| 3. Conscientiousness | .07 / -.11 | -.16 / .08 | --- | | | | | | |
| 4. Extraversion | -.05 / -.12 | .22 / .21 | .21 / .18 | --- | | | | | |
| 5. Agreeableness | -.15 / .02 | .15 / .05 | -.18 / .17 | .33* / .36* | --- | | | | |
| 6. Neuroticism | .26* / .34* | -.10 / .01 | -.42** / -.29* | -.27*/ .40** | -.14 / -.34* | --- | | | |
| 7. Trait-Anxiety | .15 / .27* | .05 / .16 | -.15 / -.22 | .20 / .01 | -.20 / -.13 | .16 / .40** | --- | | |
| 8. Intelligence | -.08 / .15 | .14 / -.09 | -.03 / -.06 | -.09 / .17 | .15 / .22 | -.21 / -.19 | -.20 / -.12 | --- | |
| 9. $HR_{change}$ | .05 / -.01 | -.11 / .04 | .02 / -.18 | -.07 / -.04 | -.21 / .09 | .02 / .10 | -.05 / -.05 | .10 / -.02 | --- |
| 10. $HR_{abs}$ | .10 / .13 | .08 / .08 | -.13 / -.30* | -.02 / -.07 | .06 / -.04 | .30* / .08 | .14 /.19 | -.17 / .20 | .03 / .26* |
| 11. $BP_{change}$ | -.13 / .02 | .00 / .05 | -.02 / .11 | -.06 / -.02 | -.08 / -.01 | .01 / -.22 | -.32* / -.08 | .20 / .07 | .28* / .07 |
| 12. $BP_{abs}$ | .08 / -.09 | .16 / .03 | .11 / -.11 | .07 /-.13 | -.22 / .01 | .01 / -.08 | -.07 / -.02 | -.04 / -.03 | .21 / .19 |
| 13. $AA_{change}$ | .03 / -.05 | -.08 / -.04 | .07 / .01 | .00 / .16 | .10 / .01 | -.18 / -.19 | -.15 / .10 | .12 / .11 | -.01 / -.31* |
| 14. $AA_{abs}$ | -.04 / -.07 | .04 / .04 | .12 / .10 | -.05 / .27* | .06 / .17 | -.21 / -.11 | -.24* / .03 | .12 / .16 | -.02 / .02 |
| 15. $Cortisol_{change}$ | .13 / -.13 | .02 / .02 | .09 / .35* | -.04 / -.09 | .08 / -.19 | -.08 / -.02 | -.21 / -.15 | .07 / -.26* | .07 / -.25* |
| 16. $Cortisol_{abs}$ | .17 / .19 | .12 / .10 | .04 / -.09 | -.02 / .08 | .13 / .01 | -.03 / .06 | -.27* / -.17 | .06 / -.09 | .08 / .14 |

*Note.* The table lists Spearman's rank-order correlation coefficients ρ between all study variables. Coefficients before the slash (/) represent correlations in the MAST group ($N = 74$), while coefficients after the slash indicate correlations in the Placebo-MAST group ($N = 75$). Change scores (indicated by subscript) were calculated such that the baseline score was subtracted from the score following stress induction or Placebo, respectively (i.e., score after experimental manipulation – baseline score). * = significant at $p < .05$, ** = significant at $p < .001$ (uncorrected for multiple comparisons; for exact $p$-values see Supplementary Table S1). HR = heart rate; BP = blood pressure; AA = alpha amylase.

**Table S1 (continued)**

*Correlations between all study variables*

| Variable | 10 | 11 | 12 | 13 | 14 | 15 | 16 |
|---|---|---|---|---|---|---|---|
| 10. $HR_{abs}$ | --- | | | | | | |
| 11. $BP_{change}$ | -.10 / -.05 | --- | | | | | |
| 12. $BP_{abs}$ | .02 / .03 | .43**/.49** | --- | | | | |
| 13. $AA_{change}$ | -.07 / .13 | .09 / .00 | .18 / .00 | --- | | | |
| 14. $AA_{abs}$ | .01 / .26* | .17 / -.11 | .29* /.08 | .55**/.26* | --- | | |
| 15. $Cortisol_{change}$ | -.20 / -.25* | .25* / -.03 | .20 / -.21 | .30* / .08 | .48**/ -.05 | --- | |
| 16. $Cortisol_{abs}$ | -.14 / .16 | .21 / .07 | .29* / -.03 | .23* / -.14 | .41**/ -.09 | .78**/ .12 | --- |

*Note.* The table lists Spearman's rank-order correlation coefficients ρ between all study variables. Coefficients before the slash (/) represent correlations in the MAST group ($N = 74$), while coefficients after the slash indicate correlations in the Placebo-MAST group ($N = 75$). Change scores (indicated by subscript) were calculated such that the baseline score was subtracted from the score following stress induction or Placebo, respectively (i.e., score after experimental manipulation – baseline score). * = significant at $p < .05$, ** = significant at $p < .001$ (uncorrected for multiple comparisons; for exact *p*-values see Supplementary Table S1). HR = heart rate; BP = blood pressure; AA = alpha amylase.

**Table S2**

*Exact p-Values for correlations among all study variables*

| Variable | 1 | 2 | 3 | 4 | 5 | 6 | 7 | 8 | 9 |
|---|---|---|---|---|---|---|---|---|---|
| 1. State psychological stress | --- | | | | | | | | |
| 2. Openness | .989 /.576 | --- | | | | | | | |
| 3. Conscientiousness | .573 / .337 | .163 / .510 | --- | | | | | | |
| 4. Extraversion | .700 / .307 | .058 / .068 | .078 / .119 | --- | | | | | |
| 5. Agreeableness | .193 / .874 | .201 / .646 | .125 / .139 | .004* / .002* | --- | | | | |
| 6. Neuroticism | .028* / .003* | .393 / .963 | <.001** / .010* | .022* / <.001** | .219 / .003* | --- | | | |
| 7. Anxiety | .204 / .020* | .645 / .163 | .216 / .062 | .094 / .916 | .094 / .262 | .164 / <.001*** | --- | | |
| 8. Intelligence | .524 / .199 | .232 / .421 | .818 / .589 | .468 / .142 | .216 / .063 | .071 / .105 | .095 / .294 | --- | |
| 9. $HR_{change}$ | .659 / .909 | .359 / .745 | .857 / .119 | .545 / .757 | .079 / .452 | .852 / .405 | .662 / .664 | .407 / .832 | --- |
| 10. $HR_{abs}$ | .408 / .281 | .510 / .471 | .279 / .010* | .839 / .568 | .617 / .751 | .011* / .499 | .233 /.104 | .149 / .082 | .813 / .027* |
| 11. $BP_{change}$ | .274 / .843 | .997 / .650 | .888 / .349 | .622 / .837 | .501 / .940 | .944 / .061 | .005* / .500 | .083 / .576 | .017* / .532 |
| 12. $BP_{abs}$ | .517 / .428 | .165 / .818 | .340 / .335 | .536 /.252 | .062 / .929 | .936 / .473 | .556 / .885 | .735 / .776 | .069 / .102 |
| 13. $AA_{change}$ | .783 / .692 | .524 / .731 | .531 / .965 | .984 / .163 | .382 / .941 | .126 / .114 | .212 / .403 | .317 / .357 | .947 / .008* |
| 14. $AA_{abs}$ | .751 / .538 | .717 / .712 | .320 / .405 | .677 / .019* | .604 / .143 | .068 / .372 | .037* / .823 | .325 / .168 | .840 / .842 |
| 15. $Cortisol_{change}$ | .257 / .252 | .856 / .891 | .430 / .002* | .705 / .430 | .522 / .094 | .503 / .855 | .067 / .190 | .565 / .022* | .551 / .033* |
| 16. $Cortisol_{abs}$ | .154 / .107 | .319 / .403 | .712 / .464 | .892 / .496 | .267 / .912 | .804 / .581 | .021* / .155 | .617 / .464 | .507 / .248 |

*Note.* The table lists exact *p*-values for Spearman's rank-order correlation coefficients ρ between all study variables (for the ρ coefficients see Table 2). Values before the slash (/) represent *p*-values in the MAST group ($N = 74$), while values after the slash indicate *p*-values in the Placebo-MAST group ($N = 75$). Absolute scores (indicated by the subscript abs) represent stress scores after experimental stress induction (MAST) or time-corresponding values in the control group, respectively (Placebo-MAST). Change scores (indicated by subscript) were calculated such that the baseline score was subtracted from the score following stress induction or Placebo, respectively (i.e., score after experimental manipulation – baseline score). * = significant at $p < .05$, ** = significant at $p < .001$ (uncorrected for multiple comparisons). HR = heart rate; BP = blood pressure; AA = alpha amylase.

**Table S2 (continued)**

*Exact p-values for correlations among all study variables*

| Variable | 10 | 11 | 12 | 13 | 14 | 15 | 16 |
|---|---|---|---|---|---|---|---|
| 10. $HR_{abs}$ | --- | | | | | | |
| 11. $BP_{change}$ | .380 / .653 | --- | | | | | |
| 12. $BP_{abs}$ | .885 / .806 | <.001** / <.001** | --- | | | | |
| 13. $AA_{change}$ | .543 / .280 | .442 / .994 | .138 / .989 | --- | | | |
| 14. $AA_{abs}$ | .935 / .027* | .154 / .343 | .014* /.488 | <.001** / .024* | --- | | |
| 15. $Cortisol_{change}$ | .090 / .030* | .032* / .826 | .092 / .077 | .011* / .515 | <.001** / .699 | --- | |
| 16. $Cortisol_{abs}$ | .247 / .176 | .073 / .556 | .013* / .790 | .047* / .232 | <.001** / .457 | <.001** / .295 | --- |

*Note.* The table lists exact *p*-values for Spearman's rank-order correlation coefficients *ρ* between all study variables (for the ρ coefficients see Table 2). Coefficients before the slash (/) represent correlations in the MAST group ($N = 74$), while coefficients after the slash indicate correlations in the Placebo-MAST group ($N = 75$). Absolute scores (indicated by the subscript abs) represent stress scores after experimental stress induction (MAST) or time-corresponding values in the control group, respectively (Placebo-MAST). Change scores (indicated by subscript) were calculated such that the baseline score was subtracted from the score following stress induction or Placebo, respectively (i.e., score after experimental manipulation – baseline score). * = significant at $p < .05$, ** = significant at $p < .001$ (uncorrected for multiple comparisons). HR = heart rate; BP = blood pressure; AA = alpha amylase.

**Table S3**

*Preliminary multiple regression models on psycho-physiological correspondence (H1)*

| | Psychological stress (MAST) | | | | | Psychological stress (placebo-MAST) | | | | |
|---|---|---|---|---|---|---|---|---|---|---|
| | β (*SE*) | *p* | $p_{adj}$ | 95% CI | $R^2$ *(p)* | β (*SE*) | *p* | $p_{adj}$ | 95% CI | $R^2$ *(p)* |
| **Model 1** | | | | | .06 (.110) | | | | | .04 (.224) |
| $HR_{abs}$ | .13 (0.11) | .255 | .308 | [-.09, .35] | | .08 (0.06) | .188 | .282 | [-.04, .20] | |
| Age | -.17 (0.10) | .100 | .201 | [-.37, .03] | | -.06 (0.06) | .345 | .345 | [-.19, .07] | |
| **Model 2** | | | | | .05 (.186) | | | | | .02 (.530) |
| $BP_{abs}$ | .05 (0.10) | .613 | .736 | [-.15, .25] | | .01 (0.07) | .881 | .881 | [-.13, .16] | |
| Age | -.18 (0.10) | .081 | .161 | [-.38, .02] | | -.07 (0.06) | .263 | .395 | [-.20, .06] | |
| **Model 3** | | | | | .04 (.218) | | | | | .04 (.279) |
| $AA_{abs}$ | -.03 (0.09) | .721 | .721 | [-.22, .15] | | -.06 (0.07) | .388 | .466 | [-.21, .08] | |
| Age | -.17 (0.10) | .110 | .219 | [-.38, .04] | | -.08 (0.07) | .234 | .351 | [-.21, .05] | |
| **Model 4** | | | | | .11* (.016) | | | | | .02 (.447) |
| $Cortisol_{abs}$ | .21 (0.09) | .024* | .036* | [.03, .39] | | .10 (0.16) | .546 | .546 | [-.22, .41] | |
| Age | -.25 (0.10) | .017* | .034* | [-.46, -.05] | | -.06 (0.07) | .382 | .459 | [-.19, .07] | |
| **Model 5** | | | | | .04 (.207) | | | | | .02 (.265) |
| $HR_{change}$ | .03 (0.12) | .832 | .928 | [-.21, .27] | | .01 (0.06) | .928 | .928 | [-.11, .13] | |
| Age | -.18 (0.10) | .084 | .167 | [-.38, .02] | | -.07 (0.06) | .265 | .398 | [-.20, .06] | |
| **Model 6** | | | | | .05 (.160) | | | | | .02 (.492) |
| $BP_{change}$ | -.08 (0.11) | .456 | .548 | [-.30, .14] | | .03 (0.07) | .679 | .679 | [-.11, .17] | |
| Age | -.17 (0.10) | .095 | .190 | [-.38, .03] | | -.07 (0.06) | .271 | .406 | [-.20, .06] | |
| **Model 7** | | | | | .04 (.203) | | | | | .04 (.276) |
| $AA_{change}$ | -.05 (0.10) | .602 | .602 | [-.24, .14] | | -.07 (0.08) | .383 | .460 | [-.22, .09] | |
| Age | -.17 (0.10) | .092 | .184 | [-.38, .03] | | -.09 (0.06) | .164 | .246 | [-.22, .04] | |
| **Model 8** | | | | | .08* (.044) | | | | | .04 (.221) |
| $Cortisol_{change}$ | .17 (0.10) | .077 | .115 | [-.02, .37] | | -.18 (0.13) | .184 | .221 | [-.44, .09] | |
| Age | -.25 (0.11) | .023* | .045* | [-.47, -.04] | | -.06 (0.06) | .324 | .324 | [-.19, .06] | |

*Note*. Absolute scores (indicated by the subscript abs) represent stress scores after experimental stress induction (MAST) or time-corresponding values in the control group, respectively (Placebo-MAST). Change scores (indicated by subscript) were calculated such that the baseline score was subtracted from the score following stress induction or Placebo, respectively (i.e., score after experimental manipulation – baseline score). * = significant at $p < .05$, ** = significant at $p < .001$. HR = heart rate; BP = blood pressure; AA = alpha amylase; β = standardized regression coefficient; SE = standard error; $p_{adj}$ = Benjamini-Hochberg corrected *p*-values.

**Table S4**

*Multiple regression models testing moderators of psycho-physiological correspondence (H2)*

| | State psychological stress (MAST) | | | | | State psychological stress (placebo-MAST) | | | | |
|---|---|---|---|---|---|---|---|---|---|---|
| | β (*SE*) | *p* | $p_{adj}$ | 95% CI | $R^2$ (*p*) | β (*SE*) | *p* | $p_{adj}$ | 95% CI | $R^2$ (*p*) |
| **Model 11** | | | | | .26* (.033) | | | | | .22 (.086) |
| Anx | .20 (0.12) | .111 | .272 | [-.05, .44] | | .13 (0.14) | .380 | .464 | [-.16, .42] | |
| Anx*$HR_{abs}$ | -.08 (0.13) | .263 | .386 | [-.34, .18] | | .08 (0.07) | .130 | .278 | [-.06, .22] | |
| Anx*$BP_{abs}$ | -.24 (0.11) | .014* | .061 | [-.46, -.03] | | .10 (0.08) | .085 | .235 | [-.05, .25] | |
| Anx*$AA_{abs}$ | .17 (0.09) | .038* | .140 | [-.02, .36] | | -.16 (0.07) | .014* | .061 | [-.31, -.02] | |
| Anx*$Cortisol_{abs}$ | -.11 (0.10) | .139 | .278 | [-.31, .09] | | -.15 (0.19) | .220 | .354 | [-.53, .24] | |
| **Model 12** | | | | | .13 (.548) | | | | | .20 (.133) |
| Anx | .03 (0.14) | .821 | .903 | [-.25, .31] | | .39 (0.13) | .003* | .021* | [.14, .64] | |
| Anx*$HR_{change}$ | -.05 (0.15) | .368 | .495 | [-.34, .24] | | .06 (0.07) | .175 | .427 | [-.07, .20] | |
| Anx*$BP_{change}$ | .00 (0.14) | .500 | .611 | [-.27, .27] | | .13 (0.08) | .068 | .214 | [-.04, .30] | |
| Anx*$AA_{change}$ | .07 (0.11) | .280 | .449 | [-.16, .29] | | -.07 (0.07) | .167 | .427 | [-.21, .07] | |
| Anx*$Cortisol_{change}$ | -.08 (0.09) | .203 | .446 | [-.26, .11] | | .27 (0.16) | .050 | .184 | [-.05, .60] | |
| **Model 13** | | | | | .31* (.007) | | | | | .10 (.721) |
| O | .03 (0.11) | .753 | .828 | [-.18, .25] | | -.01 (0.13) | .928 | .928 | [-.26, .24] | |
| O*$HR_{abs}$ | -.02 (0.10) | .411 | .573 | [-.22, .18] | | .05 (0.07) | .213 | .468 | [-.08, .19] | |
| O*$BP_{abs}$ | -.01 (0.13) | .481 | .623 | [-.27, .25] | | .07 (0.08) | .194 | .468 | [-.10, .24] | |
| O*$AA_{abs}$ | .37 (0.12) | .002* | .011* | [.13, .61] | | -.11 (0.15) | .241 | .471 | [-.41, .19] | |
| O*$Cortisol_{abs}$ | -.26 (0.11) | .009* | .051 | [-.48, -.05] | | -.08 (0.16) | .310 | .487 | [-.39, .23] | |
| **Model 14** | | | | | .22 (.089) | | | | | .08 (.850) |
| O | .03 (0.12) | .773 | .810 | [-.20, .26] | | .09 (0.11) | .428 | .524 | [-.14, .31] | |
| O*$HR_{change}$ | .15 (0.17) | .187 | .479 | [-.18, .48] | | -.04 (0.10) | .327 | .479 | [-.24, .15] | |
| O*$BP_{change}$ | -.22 (0.13) | .051 | .224 | [-.48, .04] | | -.03 (0.07) | .321 | .479 | [-.17, .11] | |
| O*$AA_{change}$ | .34 (0.14) | .009* | .067 | [.06, .62] | | .05 (0.09) | .281 | .479 | [-.12, .23] | |
| O*$Cortisol_{change}$ | -.05 (0.10) | .308 | .479 | [-.26, .16] | | .03 (0.12) | .411 | .524 | [-.21, .26] | |
| **Model 15** | | | | | .23 (.078) | | | | | .13 (.513) |
| C | .04 (0.11) | .746 | .782 | [-.18, .25] | | -.15 (0.17) | .376 | .567 | [-.49, .19] | |
| C*$HR_{abs}$ | .03 (0.08) | .347 | .567 | [-.14, .20] | | .13 (0.09) | .069 | .302 | [-.04, .30] | |
| C*$BP_{abs}$ | .23 (0.11) | .021* | .150 | [.01, .45] | | -.13 (0.10) | .096 | .310 | [-.32, .07] | |
| C*$AA_{abs}$ | -.03 (0.12) | .413 | .567 | [-.27, .21] | | -.03 (0.09) | .362 | .567 | [-.21, .14] | |
| C*$Cortisol_{abs}$ | .03 (0.11) | .387 | .567 | [-.18, .24] | | -.25 (0.22) | .135 | .328 | [-.69, .20] | |
| **Model 16** | | | | | .31* (.006) | | | | | .09 (.799) |
| C | .02 (0.11) | .858 | .858 | [-.20, .24] | | .12 (0.13) | .366 | .537 | [-.15, .39] | |
| C*$HR_{change}$ | .03 (0.09) | .383 | .537 | [-.15, .21] | | .07 (0.08) | .187 | .411 | [-.09, .24] | |
| C*$BP_{change}$ | .20 (0.12) | .050 | .157 | [-.04, .43] | | -.03 (0.12) | .415 | .537 | [-.26, .21] | |
| C*$AA_{change}$ | -.32 (0.11) | .002* | .015* | [-.53, -.10] | | .03 (0.10) | .368 | .537 | [-.17, .24] | |
| C*$Cortisol_{change}$ | .29 (0.10) | .003* | .017* | [.08, .49] | | .19 (0.19) | .167 | .408 | [-.20, .57] | |
| **Model 17** | | | | | .18 (.227) | | | | | .20 (.156) |
| N | .12 (0.12) | .312 | .651 | [-.12, .36] | | .03 (0.13) | .818 | .867 | [-.24, .30] | |
| N*$HR_{abs}$ | .06 (0.12) | .622 | .805 | [-.17, .29] | | .15 (0.07) | .031* | .171 | [.01, .29] | |
| N*$BP_{abs}$ | -.10 (0.11) | .393 | .686 | [-.33, .13] | | .04 (0.09) | .659 | .805 | [-.14, .21] | |
| N*$AA_{abs}$ | -.03 (0.13) | .827 | .867 | [-.28, .23] | | -.04 (0.10) | .723 | .837 | [-.24, .17] | |
| N*$Cortisol_{abs}$ | .10 (0.12) | .411 | .686 | [-.14, .33] | | -.13 (0.17) | .437 | .686 | [-.46, .20] | |

| | State psychological stress (MAST) | | | | | State psychological stress (placebo-MAST) | | | | |
|---|---|---|---|---|---|---|---|---|---|---|
| | β (*SE*) | *p* | $p_{adj}$ | 95% CI | $R^2$ (*p*) | β (*SE*) | *p* | $p_{adj}$ | 95% CI | $R^2$ (*p*) |
| **Model 18** | | | | | .17 (.246) | | | | | .14 (.430) |
| N | .26 (0.13) | .046* | .337 | [.00, .52] | | .20 (0.13) | .114 | .551 | [-.05, .45] | |
| N*$HR_{change}$ | .07 (0.09) | .478 | .741 | [-.12, .25] | | .08 (0.10) | .410 | .709 | [-.12, .29] | |
| N*$BP_{change}$ | -.14 (0.13) | .289 | .707 | [-.40, .12] | | -.05 (0.08) | .539 | .741 | [-.20, .11] | |
| N*$AA_{change}$ | .10 (0.12) | .419 | .709 | [-.14, .34] | | .03 (0.09) | .757 | .833 | [-.16, .21] | |
| N*$Cortisol_{change}$ | -.08 (0.12) | .520 | .741 | [-.33, .17] | | .10 (0.20) | .612 | .771 | [-.30, .51] | |
| **Model 19** | | | | | .25* (.037) | | | | | .13 (.521) |
| E | -.01 (0.10) | .887 | .892 | [-.21, .19] | | .06 (0.14) | .662 | .890 | [-.21, .33] | |
| E*$HR_{abs}$ | .21 (0.12) | .082 | .224 | [-.03, .46] | | -.08 (0.07) | .274 | .547 | [-.23, .07] | |
| E*$BP_{abs}$ | .04 (0.10) | .718 | .890 | [-.17, .25] | | .03 (0.07) | .687 | .890 | [-.11, .17] | |
| E*$AA_{abs}$ | .13 (0.09) | .149 | .364 | [-.05, .31] | | -.02 (0.08) | .805 | .890 | [-.19, .15] | |
| E*$Cortisol_{abs}$ | -.23 (0.13) | .081 | .224 | [-.49, .03] | | .21 (0.17) | .227 | .500 | [-.14, .56] | |
| **Model 20** | | | | | .14 (.420) | | | | | .11 (.641) |
| E | -.06 (0.11) | .633 | .918 | [-.28, .17] | | -.19 (0.12) | .113 | .498 | [-.43, 05] | |
| E*$HR_{change}$ | -.06 (0.18) | .714 | .918 | [-.42, .29] | | -.06 (0.06) | .329 | .759 | [-.18, .06] | |
| E*$BP_{change}$ | -.11 (0.14) | .436 | .799 | [-.40, .17] | | -.02 (0.08) | .807 | .918 | [-.17, .13] | |
| E*$AA_{change}$ | -.03 (0.10) | .755 | .918 | [-.23, .17] | | -.05 (0.09) | .551 | .918 | [-.23, .12] | |
| E*$Cortisol_{change}$ | -.02 (0.12) | .878 | .918 | [-.26, .22] | | -.11 (0.12) | .367 | .759 | [-.35, .13] | |
| **Model 21** | | | | | .29* (.011) | | | | | .15 (.401) |
| A | -.22 (0.10) | .037* | .130 | [-.43, -.01] | | .20 (0.13) | .124 | .302 | [-.06, .46] | |
| A*$HR_{abs}$ | .29 (0.11) | .011* | .081 | [.07, .51] | | -.12 (0.06) | .065 | .179 | [-.24, .01] | |
| A*$BP_{abs}$ | .14 (0.11) | .230 | .380 | [-.09, .37] | | .09 (0.08) | .242 | .380 | [-.06, .24] | |
| A*$AA_{abs}$ | .14 (0.12) | .230 | .380 | [-.09, .37] | | .06 (0.10) | .550 | .672 | [-.14, .26] | |
| A*$Cortisol_{abs}$ | -.05 (0.11) | .672 | .778 | [-.26, .17] | | .18 (0.17) | .287 | .421 | [-.15, .51] | |
| **Model 22** | | | | | .21 (.125) | | | | | .07 (.891) |
| A | -.08 (0.12) | .508 | .840 | [-.33, .16] | | .06 (0.13) | .672 | .840 | [-.21, .33] | |
| A*$HR_{change}$ | .25 (0.14) | .079 | .348 | [-.03, .54] | | -.08 (0.08) | .314 | .645 | [-.24, .08] | |
| A*$BP_{change}$ | -.19 (0.13) | .155 | .567 | [-.46, .07] | | .03 (0.07) | .640 | .840 | [-.11, .17] | |
| A*$AA_{change}$ | .06 (0.11) | .592 | .840 | [-.15, .27] | | .01 (0.08) | .861 | .947 | [-.15, .18] | |
| A*$Cortisol_{change}$ | .07 (0.11) | .535 | .840 | [-.15, .28] | | .01 (0.16) | .973 | .973 | [-.31, .32] | |
| **Model 23** | | | | | .19 (.165) | | | | | .12 (.554) |
| Int | -.06 (0.11) | .618 | .800 | [-.28, .17] | | .12 (0.13) | .368 | .713 | [-.14, .38] | |
| Int*$HR_{abs}$ | -.10 (0.13) | .455 | .714 | [-.36, .16] | | -.07 (0.08) | .409 | .713 | [-.23, .10] | |
| Int*$BP_{abs}$ | .04 (0.12) | .752 | .808 | [-.20, .28] | | .07 (0.09) | .421 | .713 | [-.10, .24] | |
| Int*$AA_{abs}$ | -.02 (0.10) | .882 | .882 | [-.22, .19] | | -.03 (0.09) | .757 | .808 | [-.20, .14] | |
| Int*$Cortisol_{abs}$ | .14 (0.09) | .107 | .394 | [-.03, .32] | | .11 (0.18) | .550 | .800 | [-.25, .46] | |
| **Model 24** | | | | | .17 (.288) | | | | | .11 (.657) |
| Int | -.03 (0.12) | .782 | .882 | [-.27, .21] | | .00 (0.12) | .980 | .980 | [-.24, .25] | |
| Int*$HR_{change}$ | .11 (0.14) | .431 | .675 | [-.17, .39] | | .06 (0.08) | .460 | .675 | [-.09, .21] | |
| Int*$BP_{change}$ | -.07 (0.14) | .632 | .818 | [-.36, .22] | | .14 (0.09) | .136 | .580 | [-.05, .33] | |
| Int*$AA_{change}$ | .14 (0.11) | .226 | .580 | [-.09, .36] | | -.08 (0.08) | .317 | .580 | [-.24, .08] | |
| Int*$Cortisol_{change}$ | .12 (0.14) | .411 | .675 | [-.17, .40] | | -.05 (0.19) | .802 | .882 | [-.42, .32] | |

**Table S5**

*Moderated regressions' main effects of absolute or change values in physiological variables and age*

| | State psychological stress (MAST) | | | | | State psychological stress (placebo-MAST) | | | | |
|---|---|---|---|---|---|---|---|---|---|---|
| | β (*SE*) | *p* | $p_{adj}$ | 95% CI | $R^2$ *(p)* | β (*SE*) | *p* | $p_{adj}$ | 95% CI | $R^2$ *(p)* |
| **Model 11** | | | | | .26* (.033) | | | | | .22 (.086) |
| $HR_{abs}$ | .14 (0.11) | .190 | .349 | [-.07, .36] | | .07 (0.06) | .225 | .354 | [-.05, .20] | |
| $BP_{abs}$ | .05 (0.10) | .646 | .677 | [-.16, .25] | | .07 (0.07) | .359 | .464 | [-.08, .21] | |
| $AA_{abs}$ | -.05 (0.10) | .605 | .666 | [-.26, .15] | | -.06 (0.08) | .414 | .479 | [-.22, .09] | |
| $Cortisol_{abs}$ | .26 (0.10) | .012* | .061 | [.06, .46] | | .06 (0.16) | .714 | .714 | [-.26, .38] | |
| Age | -.19 (0.11) | .084 | .235 | [-.41, .03] | | -.07 (0.07) | .332 | .456 | [-.21, .07] | |
| **Model 12** | | | | | .13 (.548) | | | | | .20 (.133) |
| $HR_{change}$ | .04 (0.13) | .763 | .883 | [-.22, .30] | | -.01 (0.06) | .930 | .930 | [-.12, .11] | |
| $BP_{change}$ | -.12 (0.12) | .324 | .475 | [-.37, .12] | | .01 (0.07) | .866 | .908 | [-.13, .16] | |
| $AA_{change}$ | -.09 (0.10) | .383 | .495 | [-.30, .12] | | -.09 (0.08) | .277 | .449 | [-.26, .08] | |
| $Cortisol_{change}$ | .23 (0.11) | .035* | .152 | [.02, .45] | | -.15 (0.14) | .286 | .449 | [-.42, .13] | |
| Age | -.28 (0.12) | .026* | .143 | [-.52, -.03] | | -.08 (0.06) | .225 | .449 | [-.21, .05] | |
| **Model 13** | | | | | .31 (.007) | | | | | .10 (.721) |
| $HR_{abs}$ | .24 (0.10) | .026* | .113 | [.03, .45] | | .10 (0.07) | .128 | .353 | [-.03, .24] | |
| $BP_{abs}$ | .04 (0.10) | .726 | .828 | [-.17, .24] | | .01 (0.08) | .865 | .906 | [-.14, .17] | |
| $AA_{abs}$ | -.10 (0.09) | .305 | .487 | [-.28, .09] | | -.10 (0.09) | .257 | .471 | [-.28, .08] | |
| $Cortisol_{abs}$ | .19 (0.10) | .052 | .163 | [.00, .38] | | .08 (0.18) | .654 | .788 | [-.27, .43] | |
| Age | -.22 (0.10) | .032* | .116 | [-.42, -.02] | | -.06 (0.07) | .417 | .573 | [-.21, .09] | |
| **Model 14** | | | | | .22 (.089) | | | | | .08 (.850) |
| $HR_{change}$ | .09 (0.13) | .498 | .548 | [-.17, .34] | | .01 (0.08) | .941 | .941 | [-.16, .17] | |
| $BP_{change}$ | -.17 (0.12) | .154 | .479 | [-.40, .06] | | .06 (0.09) | .483 | .548 | [-.11, .23] | |
| $AA_{change}$ | -.11 (0.10) | .269 | .479 | [-.30, .09] | | -.08 (0.09) | .358 | .492 | [-.26, .10] | |
| $Cortisol_{change}$ | .18 (0.10) | .086 | .314 | [-.03, .38] | | -.16 (0.14) | .285 | .479 | [-.44, .13] | |
| Age | -.27 (0.11) | .013* | .073 | [-.49, -.06] | | -.07 (0.07) | .319 | .479 | [-.21, .07] | |
| **Model 15** | | | | | .23 (.078) | | | | | .13 (.513) |
| $HR_{abs}$ | .16 (0.11) | .149 | .328 | [-.06, .39] | | .11 (0.07) | .099 | .310 | [-.02, .24] | |
| $BP_{abs}$ | .00 (0.11) | .967 | .967 | [-.21, .21] | | .04 (0.08) | .651 | .716 | [-.12, .19] | |
| $AA_{abs}$ | -.14 (0.10) | .172 | .344 | [-.34, .06] | | -.05 (0.09) | .566 | .655 | [-.22, .12] | |
| $Cortisol_{abs}$ | .23 (0.11) | .036* | .198 | [.02, .45] | | .12 (0.18) | .487 | .595 | [-.23, .48] | |
| Age | -.17 (0.11) | .124 | .328 | [-.40, .05] | | -.06 (0.07) | .456 | .590 | [-.20, .09] | |
| **Model 16** | | | | | .31* (.006) | | | | | .09 (.799) |
| $HR_{change}$ | .04 (0.11) | .709 | .821 | [-.18, .26] | | -.04 (0.08) | .596 | .728 | [-.19, .11] | |
| $BP_{change}$ | -.03 (0.11) | .803 | .858 | [-.24, .19] | | .02 (0.08) | .842 | .858 | [-.14, .17] | |
| $AA_{change}$ | -.14 (0.09) | .140 | .384 | [-.32, .05] | | -.07 (0.08) | .388 | .537 | [-.24, .09] | |
| $Cortisol_{change}$ | .20 (0.09) | .039* | .142 | [.01, .39] | | -.14 (0.17) | .406 | .537 | [-.28, .19] | |
| Age | -.24 (0.10) | .021* | .094 | [-.45, -.04] | | -.07 (0.07) | .336 | .537 | [-.21, .07] | |
| **Model 17** | | | | | .18 (.227) | | | | | .20 (.156) |
| $HR_{abs}$ | .12 (0.11) | .326 | .651 | [-.12, .35] | | .08 (0.06) | .222 | .651 | [-.05, .21] | |
| $BP_{abs}$ | -.01 (0.11) | .902 | .902 | [-.23, .20] | | .09 (0.07) | .254 | .651 | [-.06, .23] | |
| $AA_{abs}$ | -.11 (0.10) | .303 | .651 | [-.31, .10] | | -.06 (0.09) | .519 | .761 | [-.24, .12] | |
| $Cortisol_{abs}$ | .24 (0.10) | .025* | .171 | [.03, .45] | | .08 (0.16) | .647 | .805 | [-.25, .40] | |
| Age | .16 (0.11) | .152 | .651 | [-.39, .06] | | -.08 (0.07) | .234 | .651 | [-.22, .06] | |
| **Model 18** | | | | | .17 (.246) | | | | | .14 (.430) |
| $HR_{change}$ | .06 (0.12) | .631 | .771 | [-.19, .31] | | -.02 (0.06) | .729 | .833 | [-.15, .11] | |
| $BP_{change}$ | -.13 (0.11) | .271 | .707 | [-.36, .10] | | .07 (0.08) | .356 | .709 | [-.08, .23] | |
| $AA_{change}$ | -.01 (0.11) | .906 | .949 | [-.24, .21] | | .00 (0.09) | .955 | .955 | [-.18, .17] | |

| | State psychological stress (MAST) | | | | | State psychological stress (placebo-MAST) | | | | |
|---|---|---|---|---|---|---|---|---|---|---|
| | β (*SE*) | *p* | $p_{adj}$ | 95% CI | $R^2$ (*p*) | β (*SE*) | *p* | $p_{adj}$ | 95% CI | $R^2$ (*p*) |
| Cortisol$_{change}$ | .17 (0.11) | .132 | .551 | [-.05, .39] | | -.18 (0.14) | .208 | .654 | [-.45, .10] | |
| Age | -.17 (0.12) | .150 | .551 | [-.41, .06] | | -.06 (0.07) | .366 | .709 | [-.20, .07] | |
| **Model 19** | | | | | .25* (.037) | | | | | .13 (.521) |
| HR$_{abs}$ | .19 (0.11) | .076 | .224 | [-.02, .41] | | .14 (0.07) | .067 | .224 | [-.01, .29] | |
| BP$_{abs}$ | .01 (0.10) | .892 | .892 | [-.19, .22] | | -.05 (0.09) | .601 | .890 | [-.22, .13] | |
| AA$_{abs}$ | -.08 (0.10) | .408 | .691 | [-.28, .12] | | -.02 (0.09) | .786 | .890 | [-.20, .15] | |
| Cortisol$_{abs}$ | .19 (0.10) | .073 | .224 | [-.02, .39] | | -.05 (0.19) | .810 | .890 | [-.42, .33] | |
| Age | -.22 (0.10) | .038* | .224 | [-.43, -.01] | | -.08 (0.07) | .306 | .562 | [-.23, .07] | |
| **Model 20** | | | | | .14 (.420) | | | | | .11 (.641) |
| HR$_{change}$ | .03 (0.14) | .816 | .918 | [-.25, 32] | | -.01 (0.07) | .918 | .918 | [-.14, .12] | |
| BP$_{change}$ | -.13 (0.13) | .309 | .759 | [-.39, .12] | | .01 (0.08) | .885 | .918 | [-.15, .17] | |
| AA$_{change}$ | -.09 (0.11) | .380 | .759 | [-.30, .12] | | -.04 (0.09) | .702 | .918 | [-.22, .15] | |
| Cortisol$_{change}$ | .21 (0.11) | .063 | .348 | [-.01, .43] | | -.15 (0.16) | .345 | .759 | [-.46, .16] | |
| Age | -.22 (0.11) | .061 | .348 | [-.44, .01] | | -.10 (0.07) | .153 | .563 | [-.24, .04] | |
| **Model 21** | | | | | .29* (.011) | | | | | .15 (.401) |
| HR$_{abs}$ | .13 (0.10) | .233 | .380 | [-.08, .34] | | .17 (0.07) | .018* | .100 | [.03, .32] | |
| BP$_{abs}$ | -.03 (0.10) | .790 | .869 | [-.23, .18] | | .00 (0.08) | .980 | .980 | [-.15, .16] | |
| AA$_{abs}$ | -.08 (0.09) | .378 | .519 | [-.27, .10] | | -.13 (0.10) | .203 | .380 | [-.33, .07] | |
| Cortisol$_{abs}$ | .22 (0.10) | .031* | .130 | [.02, .41] | | .01 (0.16) | .959 | .980 | [-.32, .34] | |
| Age | -.21 (0.10) | .042* | .130 | [-.42, -.01] | | -.06 (0.07) | .420 | .543 | [-.20, .08] | |
| **Model 22** | | | | | .21 (.125) | | | | | .07 (.891) |
| HR$_{change}$ | .05 (0.13) | .688 | .840 | [-.21, .31] | | .02 (0.13) | .672 | .946 | [-.12, .15] | |
| BP$_{change}$ | -.11 (0.11) | .348 | .645 | [-.34, .12] | | .00 (0.08) | .314 | .973 | [-.16, .17] | |
| AA$_{change}$ | -.11 (0.10) | .294 | .645 | [-.31, .09] | | -.08 (0.07) | .640 | .645 | [-.25, .09] | |
| Cortisol$_{change}$ | .23 (0.11) | .045* | .245 | [.01, .45] | | -.16 (0.08) | .861 | .645 | [-.47, .16] | |
| Age | -.29 (0.11) | .011* | .079 | [-.51, -.07] | | -.09 (0.16) | .973 | .624 | [-.23, .05] | |
| **Model 23** | | | | | .19 (.165) | | | | | .12 (.554) |
| HR$_{abs}$ | .15 (0.11) | .203 | .637 | [-.08, .37] | | .11 (0.07) | .089 | .391 | [-.02, .25] | |
| BP$_{abs}$ | -.05 (0.12) | .675 | .808 | [-.28, .18] | | .02 (0.08) | .771 | .808 | [-.13, .17] | |
| AA$_{abs}$ | -.11 (0.10) | .269 | .713 | [-.31, .09] | | -.07 (0.08) | .357 | .713 | [-.23, .08] | |
| Cortisol$_{abs}$ | .27 (0.11) | .013* | .097 | [.06, .48] | | .09 (0.18) | .600 | .800 | [-.26, .45] | |
| Age | -.21 (0.11) | .054 | .295 | [-.43, .00] | | -.06 (0.07) | .396 | .713 | [-.21, .08] | |
| **Model 24** | | | | | .17 (.288) | | | | | .11 (.657) |
| HR$_{change}$ | .01 (0.13) | -964 | .980 | [-.25, .26] | | -.04 (0.08) | .599 | .818 | [-.20, .12] | |
| BP$_{change}$ | -.12 (0.12) | .305 | .580 | [-.36, .11] | | .03 (0.07) | .711 | .869 | [-.12, .18] | |
| AA$_{change}$ | -.13 (0.11) | .220 | .580 | [-.35, .08] | | -.09 (0.08) | .281 | .580 | [-.25, .07] | |
| Cortisol$_{change}$ | .19 (0.11) | .084 | .459 | [-.03, .40] | | -.18 (0.15) | .238 | .580 | [-.49, .12] | |
| Age | -.24 (0.11) | .035* | .253 | [-.46, -.02] | | -.07 (0.07) | .311 | .580 | [-.21, .07] | |

*Note*. Absolute scores (indicated by the subscript abs) represent stress scores after experimental stress induction (MAST) or time-corresponding values in the control group, respectively (Placebo-MAST). Change scores (indicated by subscript) were calculated such that the baseline score was subtracted from the score following stress induction or Placebo, respectively (i.e., score after experimental manipulation – baseline score). HR = heart rate; BP = blood pressure; AA = alpha amylase; Anx = anxiety; O = openness; C = conscientiousness; N = neuroticism; E = extraversion; A = agreeableness; Int = intelligence; β = standardized regression coefficient; *SE* = standard error; $p_{adj}$ = Benjamini-Hochberg corrected *p*-values. This table shows only the main effects of the physiological variables and age from Models 11-24. The main effect of the trait of interest and the moderation terms are displayed in Table 4. * = significant at $p < .05$, ** = significant at $p < .001$

**Table S6**

*Aggregated physiological Stress scores' descriptive Statistics and correlations with study Variables*

| | MAST | | | | Placebo-MAST | | | | Welch's *t*-test | | | |
|---|---|---|---|---|---|---|---|---|---|---|---|---|
| | *M* | *SD* | Min, Max | ω | *M* | *SD* | Min, Max | ω | *t* | df | *p* | *d* |
| Aggregated physiological $stress_{abs}$ [m] | 0.43 | 0.81 | -1.06, 2.21 | .45 | -0.44 | 0.38 | -1.37, 0.72 | .12 | -8.37** | 101.88 | <.001 | 1.39 |
| Aggregated physiological $stress_{change\ m}$ | 0.53 | 0.82 | -1.39, 2.28 | .29 | -0.53 | 0.41 | -1.62, 0.15 | .29 | -9.83** | 106.02 | <.001 | 1.63 |

*Note.* [m] = psychological and physiological variables that capture the stress reaction following the experimental manipulation in the MAST group and are thus assumed to differ between groups (i.e., manipulation check). Statistical significance of group differences was assessed with Welch's *t*-tests. For variables assumed to differ between groups, Welch's *t*-tests were conducted one-sided, while all other variables were tested two-sided. Change scores (indicated by subscript) were calculated such that the baseline score was subtracted from the score following stress induction or Placebo, respectively (i.e., score after experimental manipulation – baseline score), that is, a positive value indicates an increase in stress following experimental manipulation, while a negative value indicates a decrease in stress. Sample size (*N*) of the MAST group = 74; *N* of the Placebo-MAST group = 75. * = significant at $p < .05$, ** = significant at $p < .001$ (uncorrected for multiple comparisons).

**Table S7**

*Correlations between aggregated physiological stress scores and study variables*

| Variable | Aggregated physiological stress$_{abs}$ ρ (*p*) | | Aggregated physiological stress$_{change}$ ρ (*p*) | |
|---|---|---|---|---|
| | MAST | Placebo-MAST | MAST | Placebo-MAST |
| State psychological stress | .10 (.387) | .00 (.982) | .09 (.459) | -.20 (.090) |
| Openness | .12 (.325) | .04 (.755) | .00 (.987) | .06 (.620) |
| Conscientiousness | .14 (.244) | -.04 (.720) | .10 (.407) | .41** (<.001) |
| Extraversion | -.02 (.855) | .07 (.582) | -.05 (.693) | .01 (.935) |
| Agreeableness | .01 (.925) | .10 (.396) | .07 (.533) | -.12 (.302) |
| Neuroticism | -.09 (.460) | -.13 (.259) | -.10 (.419) | -.16 (.168) |
| Anxiety | -.31* (.008) | -.12 (.296) | -.27* (.021) | -.06 (.595) |
| Intelligence | .04 (.713) | -.08 (.495) | .11 (.371) | -.22 (.065) |
| $HR_{change}$ | .14 (.234) | .10 (.411) | .06 (.619) | -.38* (.001) |
| $HR_{abs}$ | -.15 (.216) | .08 (.507) | -.18 (.130) | -.23 (.054) |
| $BP_{change}$ | .31* (.006) | .33* (.004) | .36* (.002) | .31* (.008) |
| $BP_{abs}$ | .51** (<.001) | .61** (<.001) | .25* (.031) | .03 (.791) |
| $AA_{change}$ | .33* (.004) | .06 (<.631) | .47** (<.001) | .41** (<.001) |
| $AA_{abs}$ | .60** (<.001) | .33* (.004) | .57** (<.001) | .06 (.608) |
| $Cortisol_{change}$ | .77** (<.001) | .01 (.901) | .96** (<.001) | .80** (<.001) |
| $Cortisol_{abs}$ | .93** (<.001) | .56** (<.001) | .76** (<.001) | -.03 (.822) |

*Note.* The table lists Spearman's rank-order correlation coefficients ρ and exact *p*-values for correlations between aggregated physiological stress scores and all other study variables. *N* MAST group = 74; *N* Placebo-MAST group = 75. Absolute scores (indicated by the subscript abs) represent stress scores after experimental stress induction (MAST) or time-corresponding values in the control group, respectively (Placebo-MAST). Change scores (indicated by subscript) were calculated such that the baseline score was subtracted from the score following stress induction or Placebo, respectively (i.e., score after experimental manipulation – baseline score). * = significant at $p < .05$, ** = significant at $p < .001$ (uncorrected for multiple comparisons). HR = heart rate; BP = blood pressure; AA = alpha amylase.

**Table S8**

*Multiple regression models on psycho-physiological correspondence using aggregated physiological stress*

| | Psychological stress (MAST) | | | | | Psychological stress (placebo-MAST) | | | | |
|---|---|---|---|---|---|---|---|---|---|---|
| | β (*SE*) | *p* | $p_{adj}$ | 95% CI | $R^2$ *(p)* | β (*SE*) | *p* | $p_{adj}$ | 95% CI | $R^2$ *(p)* |
| *Preliminary Models* | | | | | | | | | | |
| **Model 25** | | | | | .07 (.071) | | | | | .03 (.403) |
| Phys. $stress_{abs}$ | .19 (0.12) | .126 | .189 | [-.05, .43] | | -.02 (0.18) | .915 | .915 | [-.38, .34] | |
| Age | -.22 (0.10) | .038* | .076 | [-.43, -.01] | | -.09 (0.07) | .179 | .215 | [-.22, .04] | |
| **Model 26** | | | | | .06 (.115) | | | | | .04 (.204) |
| Phys. $stress_{change}$ | .15 (0.12) | .237 | .243 | [-.10, .40] | | -.19 (0.16) | .243 | .243 | [-.52, .13] | |
| Age | -.22 (0.11) | .045* | .090 | [-.43, .00] | | -.08 (0.06) | .201 | .243 | [-.21, .05] | |
| *Moderation Models* | | | | | | | | | | |
| **Model 27** | | | | | .10 (.142) | | | | | .11 (.100) |
| Phys. $stress_{abs}$ | .23 (0.13) | .081 | .203 | [-.03, .48] | | .04 (0.18) | .806 | .806 | [-.31, .40] | |
| Age | -.23 (0.11) | .029* | .095 | [-.45, -.03] | | -.08 (0.06) | .236 | .377 | [-.20, .05] | |
| Anx | .15 (0.12) | .233 | .377 | [-.10, .39] | | .09 (0.10) | .365 | .421 | [-.11, .29] | |
| Anx* Phys. $stress_{abs}$ | -.13 (0.11) | .264 | .377 | [-.36, .10] | | -.14 (0.16) | .379 | .421 | [-.47, .18] | |
| **Model 28** | | | | | .07 (.314) | | | | | .13 (.055) |
| Phys. $stress_{change}$ | .16 (0.13) | .205 | .306 | [-.09, .42] | | -.21 (0.16) | .211 | .306 | [-.53, .12] | |
| Age | -.22 (0.11) | .049* | .122 | [-.43, .00] | | -.08 (0.06) | .214 | .306 | [-.20, .05] | |
| Anx | .08 (0.12) | .496 | .551 | [-.16, .33] | | .23 (0.10) | .023* | .076 | [.03, .43] | |
| Anx* Phys. $stress_{change}$ | -.05 (0.11) | .611 | .611 | [-.27, .16] | | .17 (0.16) | .284 | .355 | [-.14, .48] | |
| **Model 29** | | | | | .10 (.124) | | | | | .04 (.635) |
| Phys. $stress_{abs}$ | .19 (0.12) | .129 | .322 | [-.06, .43] | | -.03 (0.18) | .870 | .924 | [-.39, .33] | |
| Age | -.21 (0.10) | .045* | .151 | [-.42, .00] | | -.09 (0.07) | .193 | .364 | [-.22, .05] | |
| O | -.02 (0.11) | .862 | .924 | [-.23, .19] | | .07 (0.11) | .558 | .798 | [-.16, .29] | |
| O* Phys. $stress_{abs}$ | -.17 (0.14) | .219 | .364 | [-.45, .10] | | .02 (0.18) | .924 | .924 | [-.35, .38] | |
| **Model 30** | | | | | .07 (.300) | | | | | .06 (.391) |
| Phys. $stress_{change}$ | .14 (0.13) | .277 | .464 | [-.10, .40] | | -.19 (0.17) | .256 | .461 | [-.52, .14] | |
| Age | -.21 (0.11) | .051 | .169 | [-.43, .00] | | -.08 (0.07) | .204 | .461 | [-.22, .05] | |
| O | -.04 (0.10) | .722 | .724 | [-.25, .15] | | .09 (0.11) | .401 | .572 | [-.12, .30] | |
| O* Phys. $stress_{change}$ | -.07 (0.13) | .583 | .724 | [-.17, .17] | | .05 (0.14) | .724 | .724 | [-.23, .33] | |
| **Model 31** | | | | | .13* (.042) | | | | | .06 (.394) |
| Phys. $stress_{abs}$ | .15 (0.12) | .220 | .315 | [-.09, .39] | | .09 (0.19) | .649 | .721 | [-.30, .47] | |
| Age | -.21 (0.10) | .050 | .136 | [-.42, .00] | | -.07 (0.07) | .280 | .350 | [-.20, .06] | |
| C | .01 (0.10) | .902 | .902 | [-.19, .21] | | -.19 (0.13) | .152 | .254 | [-.46, .07] | |
| C* Phys. $stress_{abs}$ | .24 (0.12) | .055 | .136 | [.00, .49] | | -.39 (0.26) | .146 | .254 | [-.91, .14] | |
| **Model 32** | | | | | .12 (.066) | | | | | .06 (.372) |
| Phys. $stress_{change}$ | .13 (0.12) | .293 | .421 | [-.11, .37] | | -.16 (0.18) | .395 | .439 | [-.52, .21] | |
| Age | -.23 (0.11) | .034* | .112 | [-.44, -.02] | | -.07 (0.07) | .279 | .421 | [-.21, .05] | |
| C | -.02 (0.11) | .864 | .864 | [-.24, .20] | | .11 (0.12) | .385 | .439 | [-.07, .19] | |
| C* Phys. $stress_{change}$ | .27 (0.14) | .071 | .176 | [-.02, .55] | | .20 (0.19) | .295 | .421 | [-.17, .30] | |
| **Model 33** | | | | | .12 (.074) | | | | | .11 (.099) |
| Phys. $stress_{abs}$ | .19 (0.12) | .122 | .268 | [-.05, .43] | | .05 (0.18) | .798 | .886 | [-.31, .40] | |
| Age | -.16 (0.11) | .157 | .268 | [-.37, .06] | | -.09 (0.06) | .161 | .268 | [-.22, .04] | |
| N | .19 (0.11) | .084 | .268 | [-.03, .41] | | .07 (0.11) | .527 | .659 | [-.15, .29] | |
| N* Phys. $stress_{abs}$ | -.02 (0.12) | .898 | .898 | [-.26, .23] | | -.17 (0.18) | .339 | .485 | [-.53, .19] | |
| **Model 34** | | | | | .11 (.100) | | | | | .11 (.101) |

| | | | | | | | | | | |
|---|---|---|---|---|---|---|---|---|---|---|
| Phys. stress$_{change}$ | .16 (0.13) | .218 | .314 | [-.10, .41] | | -.16 (0.17) | .341 | .426 | [-.50, .17] | |
| Age | -.16 (0.11) | .167 | .314 | [-.38, .07] | | -.08 (0.06) | .220 | .314 | [-.21, .05] | |
| N | .22 (0.12) | .068 | .227 | [-.02, .46] | | .17 (0.11) | .124 | .311 | [-.05, .39] | |
| N* Phys. stress$_{change}$ | -.09 (0.13) | .499 | .554 | [-.34, .16] | | .05 (0.18) | .763 | .763 | [-.30, .41] | |
| **Model 35** | | | | | .10 (.116) | | | | | .06 (.345) |
| Phys. stress$_{abs}$ | .12 (0.14) | .399 | .664 | [-.16, .39] | | -.04 (0.21) | .859 | .859 | [-.45, .38] | |
| Age | -.21 (0.11) | .057 | .189 | [-.42, .01] | | -.10 (0.07) | .141 | .353 | [-.23, .03] | |
| E | -.04 (0.10) | .666 | .779 | [-.25, .16] | | -.07 (0.11) | .528 | .754 | [-.29, .15] | |
| E* Phys. stress$_{abs}$ | -.15 (0.13) | .227 | .453 | [-.40, .10] | | .08 (0.20) | .701 | .779 | [-.32, .47] | |
| **Model 36** | | | | | .08 (.206) | | | | | .09 (.179) |
| Phys. stress$_{change}$ | .11 (0.13) | .406 | .507 | [-.15, .37] | | -.16 (0.17) | .338 | .482 | [-.50, .17] | |
| Age | -.20 (0.11) | .066 | .220 | [-.42, .01] | | -.10 (0.06) | .137 | .273 | [-.23, .03] | |
| E | -.04 (0.11) | .691 | .691 | [-.25, .17] | | -.17 (0.11) | .132 | .273 | [-.39, .05] | |
| E* Phys. stress$_{change}$ | -.12 (0.12) | .321 | .482 | [-.37, .12] | | -.10 (0.14) | .478 | .531 | [-.38, .18] | |
| **Model 37** | | | | | .10 (.109) | | | | | .04 (.582) |
| Phys. stress$_{abs}$ | .19 (0.13) | .134 | .244 | [-.06, .44] | | -.07 (0.19) | .706 | .772 | [-.45, .30] | |
| Age | -.22 (0.10) | .041* | .135 | [-.42, .01] | | -.10 (0.07) | .146 | .244 | [-.23, .03] | |
| A | -.15 (0.10) | .136 | .244 | [-.36, .05] | | .12 (0.12) | .324 | .425 | [-.12, .35] | |
| A* Phys. stress$_{abs}$ | .03 (0.12) | .772 | .772 | [-.20, .27] | | .21 (0.22) | .340 | .425 | [-.23, .66] | |
| **Model 38** | | | | | | | | | | .05 (.507) |
| Phys. stress$_{change}$ | .16 (0.13) | .215 | .349 | [-.09, .41] | | -.22 (0.19) | .244 | .349 | [-.60, .15] | |
| Age | -.22 (0.11) | .048* | .159 | [-.43, .00] | | -.09 (0.07) | .191 | .349 | [-.22, .04] | |
| A | -.17 (0.11) | .146 | .349 | [-.39, .06] | | .05 (0.13) | .684 | .767 | [-.20, .30] | |
| A* Phys. stress$_{change}$ | .03 (0.13) | .807 | .807 | [-.23, .29] | | .08 (0.19) | .690 | .767 | [-.30, .46] | |
| **Model 39** | | | | | .11 (.081) | | | | | .04 (.558) |
| Phys. stress$_{abs}$ | .16 (0.12) | .189 | .315 | [-.08, .41] | | .02 (0.19) | .899 | .899 | [-.35, .39] | |
| Age | -.19 (0.10) | .073 | .204 | [-.40, .02] | | -.09 (0.07) | .187 | .315 | [-.22, .04] | |
| Int | -.07 (0.10) | .498 | .553 | [-.27, .13] | | .12 (0.11) | .287 | .410 | [-.10, .35] | |
| Int* Phys. stress$_{abs}$ | .19 (0.11) | .082 | .204 | [-.03, .41] | | .20 (0.20) | .338 | .422 | [-.21, .60] | |
| **Model 40** | | | | | .08 (.194) | | | | | .05 (.526) |
| Phys. stress$_{change}$ | .13 (0.13) | .303 | .433 | [-.12, .38] | | -.18 (0.17) | .292 | .433 | [-.53, .16] | |
| Age | -.20 (0.11) | .070 | .234 | [-.41, .02] | | -.08 (0.07) | .203 | .405 | [-.22, .05] | |
| Int | -.07 (0.11) | .497 | .621 | [-.29, .14] | | .03 (0.11) | .823 | .914 | [-.20, .25] | |
| Int* Phys. stress$_{change}$ | .15 (0.11) | .185 | .405 | [-.07, .36] | | .02 (0.17) | .925 | .925 | [-.33, .36] | |

*Note*. Phys. stress = aggregated physiological stress. Absolute scores (indicated by the subscript abs) represent stress scores after experimental stress induction (MAST) or time-corresponding values in the control group, respectively (Placebo-MAST). Change scores (indicated by subscript) were calculated such that the baseline score was subtracted from the score following stress induction or Placebo, respectively (i.e., score after experimental manipulation – baseline score). HR = heart rate; BP = blood pressure; AA = alpha amylase; β = standardized regression coefficient; SE = standard error; $p_{adj}$ = Benjamini-Hochberg corrected *p*-value. * = significant at $p < .05$, ** = significant at $p < .001$
